# Cytosolic S100A8/A9 promotes Ca^2+^ supply at LFA-1 adhesion clusters during neutrophil recruitment

**DOI:** 10.1101/2024.04.16.589738

**Authors:** Matteo Napoli, Roland Immler, Ina Rohwedder, Valerio Lupperger, Johannes Pfabe, Mariano Gonzalez Pisfil, Anna Yevtushenko, Thomas Vogl, Johannes Roth, Melanie Salvermoser, Steffen Dietzel, Marjan Slak Rupnik, Carsten Marr, Barbara Walzog, Markus Sperandio, Monika Pruenster

**Affiliations:** Walter Brendel Center of Experimental Medicine, Biomedical Center, Institute of Cardiovascular Physiology and Pathophysiology, Ludwig-Maximilians-University, Planegg-Martinsried, Germany; Institute of AI for Health, Helmholtz Zentrum München - German Research Center for Environmental Health, Neuherberg, Germany; Center for Physiology and Pharmacology, Medical University of Vienna, Vienna, Austria; Institute of Immunology, University of Muenster, Muenster, Germany

## Abstract

S100A8/A9 is an endogenous alarmin secreted by myeloid cells during many acute and chronic inflammatory disorders. Despite increasing evidence of the proinflammatory effects of extracellular S100A8/A9, little is known about its intracellular function. Here, we show that cytosolic S100A8/A9 is indispensable for neutrophil post-arrest modifications during outside-in signaling under flow conditions in vitro and neutrophil recruitment in vivo, independent of its extracellular functions. Mechanistically, genetic deletion of S100A9 in mice (*Mrp14^−/−^*, functional *S100a8/a9^−/−^*) caused dysregulated Ca^2+^ signatures in activated neutrophils resulting in reduced Ca^2+^ availability at the formed LFA-1/F-actin clusters with defective β_2_ integrin outside-in signaling during post-arrest modifications. Consequently, we observed impaired cytoskeletal rearrangement, cell polarization and spreading, as well as cell protrusion formation in *Mrp14^−/−^* compared to WT neutrophils, making *Mrp14^−/−^* cells more susceptible to detach under flow, thereby preventing efficient neutrophil recruitment and extravasation into inflamed tissue.

**One-sentence summary:** intracellular S100A8/A9 is indispensable for firm leukocyte adhesion under flow

## INTRODUCTION

Neutrophils are the most abundant circulating leukocyte subpopulation in humans and are rapidly mobilized from the bone marrow to the circulation upon sterile inflammation and/or bacterial/viral infection [1]. The interplay between activated endothelial cells and circulating neutrophils leads to a tightly regulated series of events described as leukocyte recruitment cascade [2]. Tissue-derived proinflammatory signals provoke expression of selectins on the inflamed endothelium that capture free floating neutrophils from the bloodstream by triggering tethering and rolling through interaction with selectin ligands on the neutrophil surface [3]. Selectin mediated rolling allows neutrophils to engage with immobilized chemokines and other proinflammatory mediators such as leucotriene B4 (LTB4), N-formylmethionyl-leucyl-phenylalanine (fMLF) and various agonists for Toll-like receptors (TLRs) like TLR2, TLR4 and TLR5, presented on the endothelial surface and resulting in the activation of β_2_ integrins on neutrophils [4–7]. High affinity β_2_ integrin interaction with their corresponding receptors on the endothelium induces downstream outside-in signaling leading to post-arrest modifications such as cell spreading, adhesion strengthening and neutrophil crawling, critical requirements for successful recruitment of neutrophils into inflamed tissue [8, 9]. Accordingly, impairment in those steps favors neutrophil detachment under shear flow and re-entry of neutrophils into the blood stream [5].

Local regulation of intracellular calcium (Ca^2+^) levels is critical to synchronize rolling, arrest and polarization [8, 10]. During rolling, neutrophils show only minor Ca^2+^ activity, but a rapid increase in intracellular Ca^2+^ signaling is registered during transition from slow rolling to firm adhesion and subsequent polarization of neutrophils [11].

Neutrophil transition from rolling into firm arrest involves conformational changes of the integrin lymphocyte function-associated antigen (LFA-1) into a high affinity state allowing bond formation with intercellular adhesion molecule-1 (ICAM-1) expressed on inflamed endothelium. Tension on focal clusters of LFA-1/ICAM-1 bonds mediated by shear stress promotes the assembly of cytoskeletal adaptor proteins to integrin tails and mediates Ca^2+^-release activated (CRAC) channel ORAI-1 recruitment to focal adhesion clusters ensuring high Ca^2+^ concentrations at the “inflammatory synapse” [10]. Finally, shear stress mediated local bursts of Ca^2+^ promote assembly of the F-actin cytoskeleton allowing pseudopod formation and transendothelial migration (TEM) [8, 10, 12–14].

S100A8/A9, also known as MRP8/14 or calprotectin, is a member of the EF-hand family of proteins and the most abundant cytosolic protein complex in neutrophils [15]. Secretion of S100A8/A9 can occur via passive release of the cytosolic protein due to cellular necrosis or neutrophil extracellular trap (NET) formation [16]. Active release of S100A8/A9 without cell death can be induced by the interaction of L-selectin/PSGL-1 with E-selectin during neutrophil rolling on inflamed endothelium [6, 17, 18]. We have recently shown that E-selectin induced S100A8/A9 release occurs through a NLRP3 inflammasome dependent pathway involving GSDMD pore formation. Pore formation is a time-limited and transient process, which is reversed by the activation of the ESCRT-III machinery membrane repair mechanism [19]. Once released, the protein acts as an alarmin, exerting its proinflammatory effects on different cell types like endothelial cells, lymphocytes and neutrophils [16, 20].

In the present study, we focused on the cytosolic function of S100A8/A9 in neutrophils. We demonstrate its unique role in supplying Ca^2+^ at LFA-1 adhesion clusters during neutrophil recruitment thereby orchestrating Ca^2+^ dependent post-arrest modifications, which are critical steps for subsequent transmigration and extravasation of these cells into inflamed tissues.

## RESULTS

### Cytosolic S100A8/A9 promotes leukocyte recruitment in vivo regardless of extracellular S100A8/A9 functions

As demonstrated previously by our group, rolling of neutrophils on inflamed endothelium leads to E-selectin mediated, NLRP3 inflammasome dependent, secretion of S100A8/A9 via transient GSDMD pores [19]. Released S100A8/A9 heterodimer in turn binds to TLR4 on neutrophils in an autocrine manner, leading to β_2_ integrin activation, slow leukocyte rolling and firm neutrophil adhesion [6]. Interestingly, E-selectin-triggered S100A9/A9 release does not substantially affect the cytosolic S100A8/A9 content. Analysis of S100A8/A9 levels in the supernatants of E-selectin versus Triton X-100 treated neutrophils demonstrated that only about 1-2% of the cytosolic S100A8/A9 content was secreted to the extracellular compartment (Fig. 1A). In addition, immunofluorescence analysis of the inflamed cremaster muscle tissue confirmed no major difference in the amount of cytosolic S100A8/A9 between intravascular and extravasated neutrophils (Fig S1A and S1B). Given the abundance of cytosolic S100A8/A9 even after its active release during neutrophil rolling, we wanted to investigate a putative role of intracellular S100A8/A9 in leukocyte recruitment independently of its extracellular function.

**Figure 1:**
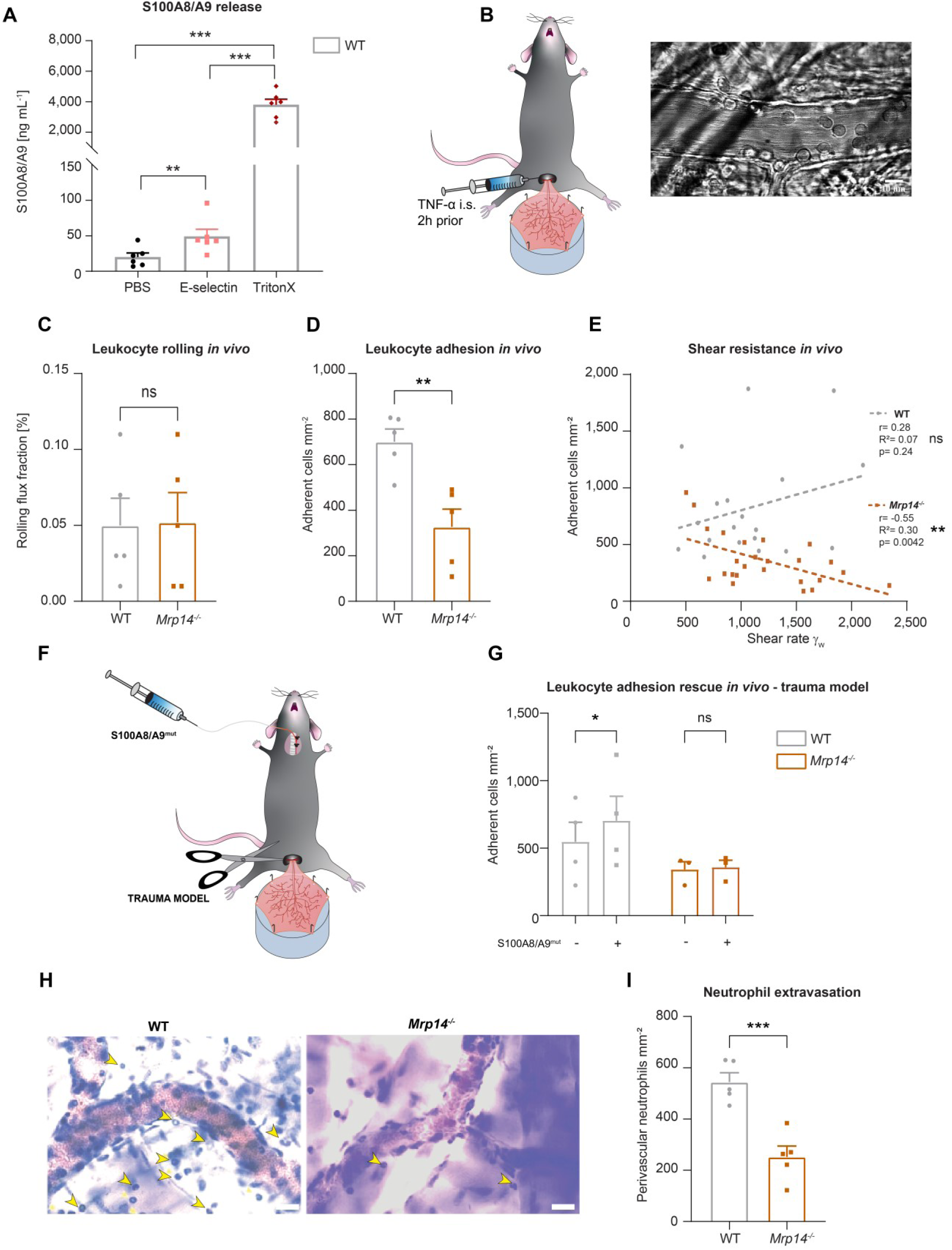
Cytosolic S100A8/A9 regulates leukocyte recruitment in vivo regardless of extracellular S100A8/A9. (A) ELISA measurements of S100A8/A9 levels in supernatants of WT bone marrow neutrophils stimulated for 10min with PBS, E-selectin or lysed with Triton X-100(mean+SEM, *n=*6 mice per group, RM one-way ANOVA, Holm-Sidak’s multiple comparison). (B) Schematic model of the mouse cremaster muscle preparation for intravital microscopy and representative picture of a vessel showing rolling and adherent cells. WT and *Mrp14^−/−^* mice were stimulated i.s. with TNF-α 2h prior to cremaster muscle post-capillary venules imaging by intravital microscopy. Quantification of (C) number or rolling (rolling flux fraction) and (D) number of adherent neutrophils per vessel surface of WT and *Mrp14^−/−^* mice [mean+SEM, *n=*5 mice per group, 25 (WT) and 30 (*Mrp14^−/−^*) vessels, unpaired Student’s *t*-test]. (E) Correlation between physiological vessel shear rates and number of adherent neutrophils in WT and *Mrp14^−/−^* mice [*n=*25 (WT) and 30 (*Mrp14^−/−^*) vessels of 5 mice per group, Pearson correlation]. (F) Schematic model of sterile inflammation induced by exteriorizing WT and *Mrp14^−/−^* cremaster muscles. (G) Analysis of number of adherent leukocytes by intravital microscopy before and after S100A8/A9^mut^ intra-arterial injection [mean+SEM, *n=*3 mice per group, 3 (WT) and 3 (*Mrp14^−/−^*) vessels, 2way ANOVA, Sidak’s multiple comparison]. (H) Representative Giemsa staining micrographs of TNF-α stimulated WT and *Mrp14^−/−^* cremaster muscles (representative micrographs, scale bar =30µm, arrows: transmigrated neutrophils) and (I) quantification of number of perivascular neutrophils [mean+SEM, *n=*5 mice per group, 56 (WT) and 55 (*Mrp14*^−/−^) vessels, unpaired Student’s *t*-test]. ns, not significant; *p≤0.05, **p≤0.01, ***p≤0.001.

To investigate this, we made use of WT and *Mrp14^−/−^*mice, which are functional double knockout mice for MRP8 and MRP14 (S100A8 and S100A9) at the protein level [21], and studied neutrophil recruitment in mouse cremaster muscle venules upon TNF-α treatment (Fig. 1B), a well-established model to assess neutrophil recruitment into inflamed tissue in vivo [22]. Two hours after onset of inflammation, we exteriorized the cremaster muscle and investigated the number of rolling and adherent cells by intravital microscopy. While rolling was not affected by the absence of S100A8/A9 (Fig. 1C), we detected a reduced number of adherent neutrophils in postcapillary cremaster muscle venules of *Mrp14^−/−^* compared to WT mice (Fig. 1D). We found a significant negative correlation between increasing shear rates and the number of adherent leukocytes in *Mrp14^−/−^*animals while this correlation could not be detected in WT mice (Fig. 1E). These findings indicate that lack of cytosolic S100A8/A9 impairs shear stress resistance of adherent neutrophils in vivo. To exclude differences in surface expression of rolling and adhesion relevant molecules on neutrophils, we performed FACS analysis and could not detect differences in the baseline expression of CD11a, CD11b, CD18, CD62L, PSGL1, CXCR2 and CD44 in WT and *Mrp14^−/−^* neutrophils (Fig. S1C - S1J). In order to test whether the observed phenotype of decreased neutrophil adhesion in *Mrp14^−/−^*mice was simply a consequence of the lack of extracellular S100A8/A9 induced β_2_ integrin activation, we again performed intravital microscopy in the exteriorized but otherwise unstimulated mouse cremaster muscles. In this scenario, only a mild inflammation is induced which leads to the mobilization of pre-stored P-selectin from Weibel-Pallade bodies, but no upregulation of E-selectin and therefore no E-selectin induced S100A8/A9 release [6]. After exteriorization and trauma-induced induction of inflammation in the cremaster muscle tissue, we systemically injected soluble S100A8/A9 via a carotid artery catheter to induce TLR4 mediated integrin activation and firm leukocyte adhesion in exteriorized cremaster muscle venules (Fig. 1F) [6]. To prevent S100A8/A9 tetramerization in plasma, which would abolish binding of S100A8/A9 to TLR4 [23], we took advantage of a mutant S100A8/A9 protein (S100A8/A9^mut^, aa exchange N70A and E79A) which is unable to tetramerize upon Ca^2+^ binding thereby inducing substantial TLR4 downstream signaling [24, 25]. Injection of S100A8/A9^mut^ induced a significant increase in leukocyte adhesion in WT mice (Fig. 1G), whereas induction of adhesion was completely absent in *Mrp14^−/−^* mice (Fig. 1G), suggesting that loss of S100A8/A9 causes an intrinsic adhesion defect, which cannot be rescued by application of extracellular S100A8/A9 and subsequent TLR4 mediated β_2_ integrin activation. In addition, similar results were obtained in the TNF-α stimulated cremaster muscles model (Fig. S1K) where S100A8/A9^mut^ increased leukocyte adhesion in WT mice, but again could not induce an increase in leukocyte adhesion in *Mrp14^−/−^* mice (Fig. S1L). In addition, microvascular parameters were quantified in order to compare different vessels in every in vivo experiment and no difference was detected (Table S1).

Further, we wanted to investigate whether reduced adhesion results in impaired leukocyte extravasation in *Mrp14^−/−^* mice and stained TNF-α stimulated cremaster muscles of WT and *Mrp14^−/−^*mice with Giemsa and analyzed number of perivascular neutrophils. Indeed, we observed a reduced number of transmigrated neutrophils in *Mrp14^−/−^*compared to WT mice (Fig. 1H and 1I). Taken together, these data indicate that cytosolic S100A8/A9 regulates key processes during neutrophil recruitment into inflamed tissue in vivo.

### Loss of cytosolic S100A8/A9 impairs neutrophil adhesion under flow conditions without affecting β_2_ integrin activation

Next, we focused on the adhesion defect of S100A8/A9 deficient neutrophils. For this purpose, we used an autoperfused microflow chamber system as described earlier [26]. Flow chambers were coated with E-selectin, ICAM-1 and CXCL1 (Fig. 2A). This combination of recombinant proteins mimics the inflamed endothelium and allows studying leukocyte adhesion under flow conditions. In line with our in vivo findings, lack of S100A8/A9 did not affect leukocyte rolling (Fig. 2B), but resulted in a lower number of adherent *Mrp14^−/−^* leukocytes compared to WT leukocytes (Fig. 2C), without affecting white blood cell count (WBC) (Table S2). Reduced neutrophil adhesion could be a consequence of defective β_2_ integrin activation induced by chemokines or other inflammatory mediators [5–7, 27]. In order to study the effect of S100A8/A9 deficiency on rapid β_2_ integrin activation via G_αi_ coupled signaling (inside-out signaling), we investigated the capacity of WT and *Mrp14^−/−^* neutrophils to bind soluble ICAM-1 upon CXCL1 stimulation using flow cytometry (Fig. 2D). CXCL1 induced a significant and similar increase in soluble ICAM-1 binding in both, WT and *Mrp14^−/−^* neutrophils (Fig. 2E), suggesting that G_αi_ coupled integrin activation is independent of cytosolic S100A8/A9. To corroborate this finding, we performed a static adhesion assay where we plated WT and *Mrp14^−/−^* neutrophils on ICAM-1 coated plates, stimulated them with PBS or CXCL1 and quantified the number of adherent cells. As expected, CXCL1 stimulated WT cells displayed increased adhesion to ICAM-1 coated plates compared to PBS treatment (Fig. 2F). In line with the findings from the soluble ICAM-1 binding assay, this increase was also detected in *Mrp14*^−/−^ cells indicating that chemokine-induced β_2_ integrin activation is not dependent on cytosolic S100A8/A9.

**Figure 2.**
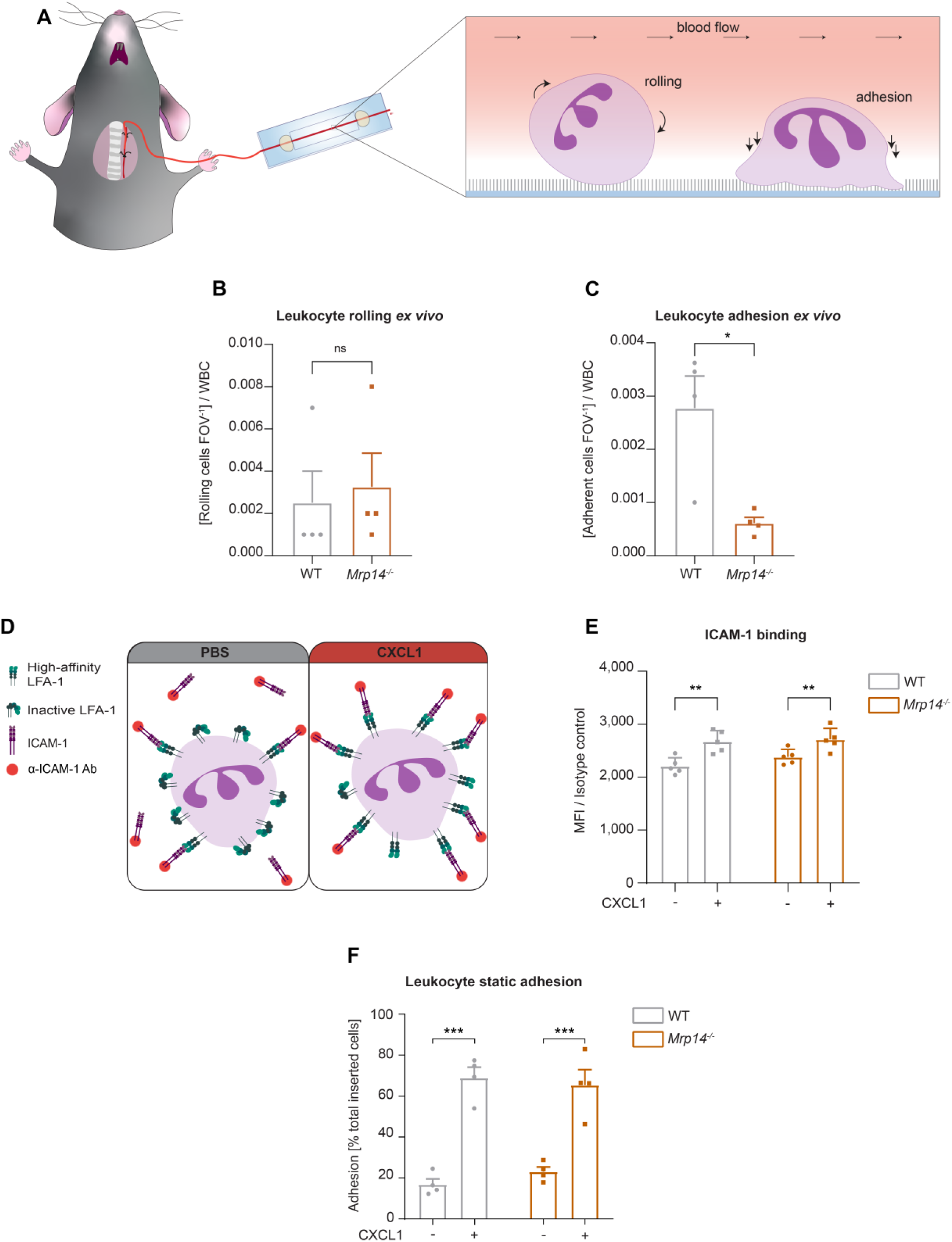
Loss of cytosolic S100A8/A9 impairs neutrophil adhesion under flow conditions without affecting β_2_ integrin activation. (A) Schematic representation of blood harvesting from WT and *Mrp14^−/−^* mice via a carotid artery catheter and perfusion into self-made flow cambers coated with E-selectin, ICAM-1, and CXCL1. Analysis of (B) number of rolling and (C) number of adherent leukocytes FOV^−1^[mean+SEM, *n=*4 mice per group, 10 (WT) and 12 (*Mrp14^−/−^*) flow chambers, paired Student’s *t*-test]. (D) Schematic representation of the soluble ICAM-1 binding assay using bone marrow neutrophils stimulated with PBS control or CXCL1 (10nM) assessed by (E) flow cytometry (MFI=median fluorescence intensity, mean+SEM, *n=*5 mice per group, 2way ANOVA, Sidak’s multiple comparison). (F) Spectroscopy fluorescence intensity analysis of percentage of adherent WT and *Mrp14^−/−^* neutrophils, seeded for 5min on ICAM-1 coated plates and stimulated with PBS or CXCL1 (10nM) for 10min (mean+SEM, *n=*4 mice per group, 2way ANOVA, Sidak’s multiple comparison). ns, not significant; *p≤0.05, **p≤0.01, ***p≤0.001.

### Cytosolic S100A8/A9 is crucial for neutrophil spreading, crawling and post-arrest modifications under flow

Activated and ligand bound β_2_ integrins start to assemble focal clusters thereby transmitting signals into the inner cell compartment [28]. This process named outside-in signaling is required to strengthen adhesion and to induce cell shape changes, fundamental for neutrophil spreading, crawling and finally transmigration [29]. Since *Mrp14^−/−^*neutrophils displayed a defect in leukocyte adhesion in vivo and ex vivo, although their inside-out signaling is fully functional, we started to study a putative role of cytosolic S100A8/A9 in β_2_ integrin dependent outside-in signaling. Therefore, isolated WT and *Mrp14^−/−^* bone marrow neutrophils were introduced into E-selectin, ICAM-1 and CXCL1 coated microflow chambers and changes in cell shape were monitored over 10min (Fig. 3A). WT neutrophils displayed normal spreading properties as depicted by the gradual increase in area and perimeter over time (Fig. 3B). In line with these findings, circularity and solidity, parameters reflecting the polarization capability of the cells and the amount of protrusions the cell developed, respectively, decreased over time (Fig. 3C). In contrast, increment of area and perimeter was significantly less pronounced in *Mrp14^−/−^* cells (Fig. 3B). Circularity and solidity did only marginally decrease over time in *Mrp14^−/−^* cells, suggesting that neutrophils are unable to polarize properly and to extend protrusions (Fig. 3C). These results imply a substantial role of cytosolic S100A8/A9 in β_2_ integrin outside-in signaling.

**Figure 3:**
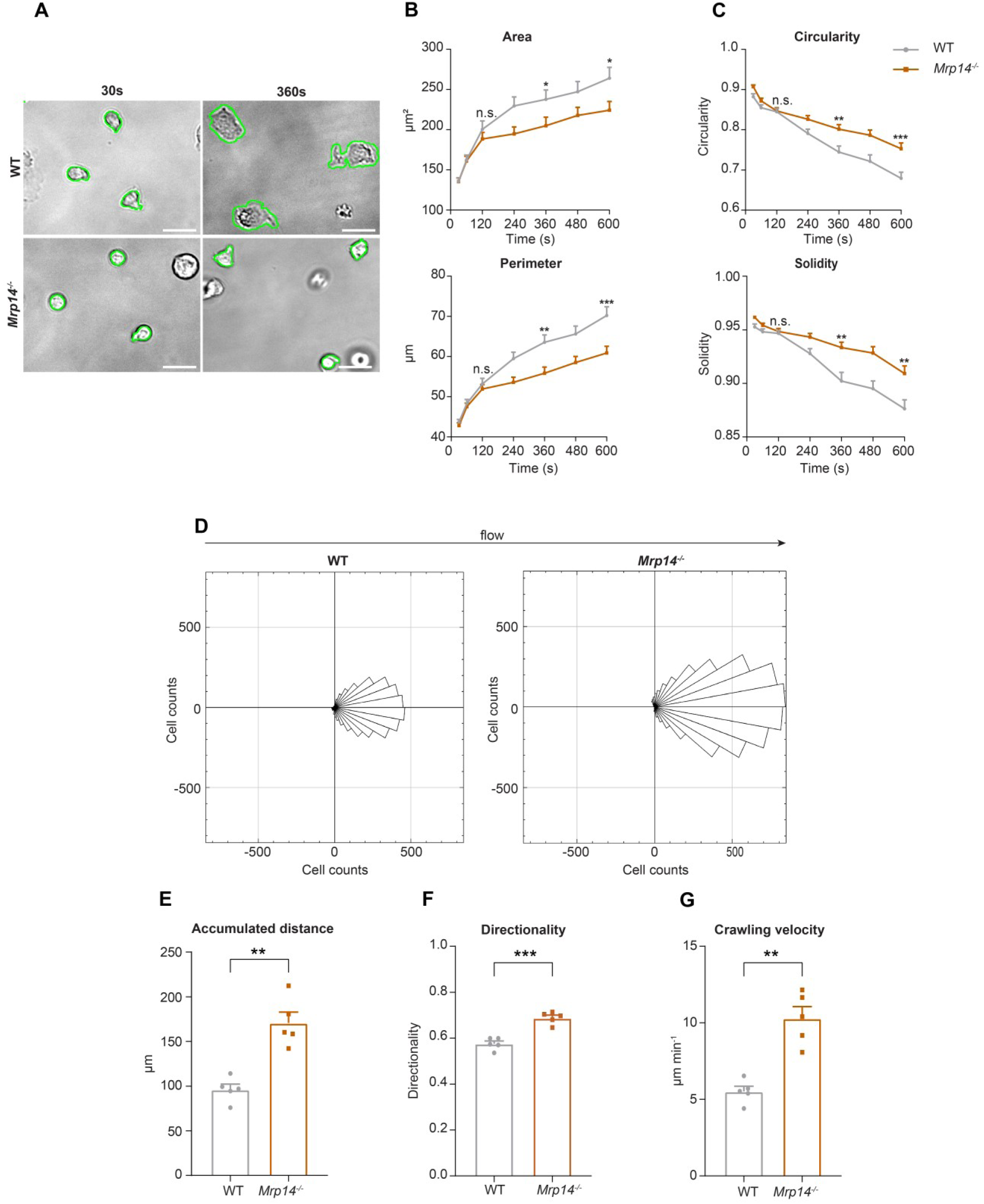
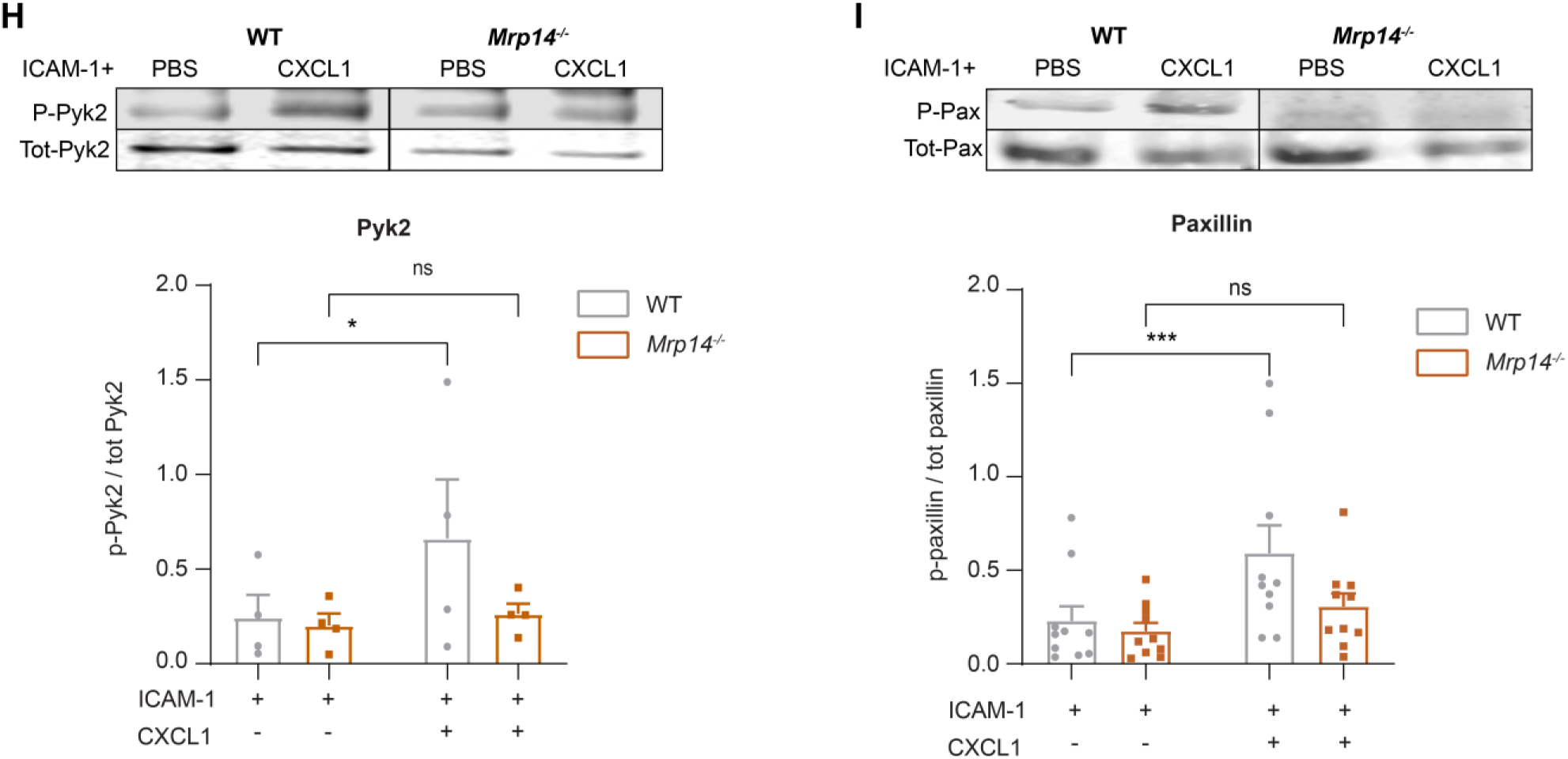
Cytosolic S100A8/A9 is crucial for neutrophil spreading, crawling and post-arrest modifications under flow. (A) Representative bright-field pictures of WT and *Mrp14^−/−^* neutrophils spreading over E-selectin, ICAM-1, and CXCL1 coated glass capillaries (scale bar=10μm). Analysis of cell shape parameters (B) area, perimeter, (C) circularity [4π * (Area/Perimeter)] and solidity [Area/Convex area] over time [mean+SEM, *n=*103 (WT) and 96 (*Mrp14^−/−^*) neutrophils of 4 mice per group, unpaired Student’s *t*-test]. (D) Rose plot diagrams representative of migratory crawling trajectories of WT and *Mrp14^−/−^* neutrophils in flow chambers coated with E-selectin, ICAM-1, and CXCL1 under flow (2dyne cm^-2^). Analysis of (E) crawling distance, (F) directionality of migration and (G) crawling velocity of WT and *Mrp14^−/−^* neutrophils [mean+SEM, *n=*5 mice per group, 113 (WT) and 109 (*Mrp14^−/−^*) cells, paired Student’s *t*-test]. Western blot analysis of ICAM-1 induced (H) Pyk2 and (I) Paxillin phosphorylation of WT and *Mrp14^−/−^* neutrophils upon CXCL1 stimulation (10nM) (mean+SEM, representative western blot of *n≥*4 mice per group, 2way ANOVA, Sidak’s multiple comparison). ns, not significant; *p≤0.05, **p≤0.01, ***p≤0.001.

Next, we wanted to examine consequences of impaired neutrophil spreading in absence of S100A8/A9 by analyzing neutrophil crawling under flow. Therefore, we introduced isolated neutrophils into E-selectin, ICAM-1 and CXCL1 coated microflow chambers and allowed them to adhere for 3min to the substrates. Thereafter, we applied physiological shear stress (2dyne cm^-2^) and analyzed crawling behavior. WT neutrophils resisted shear forces and slowly crawled in the direction of the flow, whereas *Mrp14^−/−^* neutrophils crawled in an intermittent and jerky manner (Fig. 3D and Movie S1). In line, *Mrp14^−/−^* neutrophils covered significantly longer distances (Fig. 3E), with an increased directionality toward flow direction (Fig. 3F) and displayed an increased crawling velocity compared to WT cells (Fig. 3G).

To confirm impaired crawling and defective outside-in signaling dependent adhesion strengthening in neutrophils lacking cytosolic S100A8/A9, we conducted a neutrophil detachment assay using E-selectin, ICAM-1 and CXCL1 coated microflow chambers and applied increasing shear stress. We found lower numbers of adherent *Mrp14^−/−^* neutrophils compared to WT neutrophils with increasing shear stress (Fig. S2A). This is in line with our in vivo findings where we detected a negative correlation between the number of adherent cells and increasing shear stress in *Mrp14^−/−^*animals (Fig. 1E). Together, these findings suggest a critical role of intracellular S100A8/A9 in adherent neutrophils to resist high shear stress conditions.

Following engagement of the ligand ICAM-1 to activated β_2_ integrins in neutrophils, the proline-rich tyrosine kinase Pyk2 and the focal adhesion adaptor protein paxillin are, among other proteins, rapidly tyrosine phosphorylated thereby being critical events for cell adhesion, migration and podosome formation [27, 30, 31]. To test the role of cytosolic S100A8/A9 in mediating outside-in signaling events on the mechanistic level, we seeded WT and *Mrp14^−/−^* neutrophils on ICAM-1 coated plates, stimulated the cells with CXCL1 and determined Pyk2 and paxillin phosphorylation by western blot analysis. We found increased abundance of Pyk2 and paxillin phosphorylation in CXCL1 stimulated WT cells, while no increase was detectable in *Mrp14^−/−^*neutrophils (Fig. 3H and 3I). Taken together, these data indicate that cytosolic S100A8/A9 is essential during ICAM-1 induced integrin outside-in signaling events and therefore indispensable for post arrest modifications including cell polarization and the formation of cell protrusions.

### Cytosolic S100A8/A9 drives neutrophil cytoskeletal rearrangement by regulating LFA-1 nanocluster formation and Ca^2+^ availability within the clusters

Integrin outside-in signaling strongly depends on focal cluster formation of high-affinity LFA-1 and high Ca^2+^ concentrations within these clusters [10, 12–14]. Since S100A8/A9 is a Ca^2+^ binding protein, we studied LFA-1 clustering and Ca^2+^ signatures during neutrophil adhesion under flow conditions. For this approach, we isolated neutrophils from Ca^2+^ reporter mice (WT*^Lyz2xGCaMP5^*) and S100A8/A9 deficient Ca^2+^ reporter mice (*Mrp14^−/− Lyz2xGCaMP5^*) and fluorescently labelled the cells with an LFA-1 antibody (Fig.4a). Neutrophils were then introduced into E-selectin, ICAM-1 and CXCL1 coated flow chambers, allowed to settle for 3min before shear was applied (2dyne cm^-2^). Time-lapse movies of fluorescence LFA-1 and Ca^2+^ signals were recorded for 10min by confocal microscopy. First, LFA-1 signals from single cell analysis (Fig. 4A) were segmented through automatic thresholding in order to generate a binary image of the LFA-1 signals (LFA-1 mask) (Fig. 4B). Then, LFA-1 nanoclusters were considered as such if they spanned a minimum area of 0.15µm^2^ (Fig. 4C), according to literature [32]. We found that *Mrp14^−/− Lyz2xGCaMP5^*neutrophils formed significantly less LFA-1 nanoclusters compared to WT *^Lyz2xGCaMP5^* neutrophils suggesting an involvement of cytosolic S100A8/A9 in LFA-1 nanocluster formation (Fig. 4D and Movie S2). Next, we investigated Ca^2+^ intensities within LFA-1 nanoclusters (Fig. 4E) to determine Ca^2+^ levels at the LFA-1 focal adhesion spots (Fig. 4F). We found a significant reduction of Ca^2+^ levels in LFA-1 nanocluster areas of *Mrp14^−/− Lyz2xGCaMP5^* neutrophils compared to WT *^Lyz2xGCaMP5^* neutrophils (Fig. 4G and Movie S3) suggesting an impaired availability of free intracellular Ca^2+^ at LFA-1 nanocluster sites in absence of cytosolic S100A8/A9. Strikingly, Ca^2+^ levels in the cytoplasm (outside of LFA-1 nanoclusters, Fig. 4H and 4I) did not differ between WT *^Lyz2xGCaMP5^* and *Mrp14^−/− Lyz2xGCaMP5^* neutrophils (Fig. 4J), suggesting that cytosolic S100A8/A9 plays an important role especially in supplying Ca^2+^ at LFA-1 adhesion spots. To investigate localization of S100A8/A9 during neutrophil post-arrest modification, we isolated neutrophils from WT mice and labeled them with the cell tracker green CMFDA and an LFA-1 antibody. The cells were introduced into flow chambers coated with E-selectin, ICAM-1, and CXCL1, allowed to settle for 3min, and then subjected to continuous shear stress (2dyne cm^-2^) for 10min. After fixation and permeabilization, the cells were stained for intracellular S100A9. LFA-1 nanoclusters were identified, and S100A9 intensity in these clusters was compared to that in cytoplasmic areas outside the nanoclusters. We observed higher S100A9 intensity at LFA-1 nanoclusters compared to the rest of the cytoplasm (non LFA-1 nanoclusters) in stimulated WT neutrophils (**Fig. 4K and 4L**), indicating that S100A8/A9 localizes at LFA-1 nanocluster sites, supplying them with Ca^2+^ locally.

**Figure 4:**
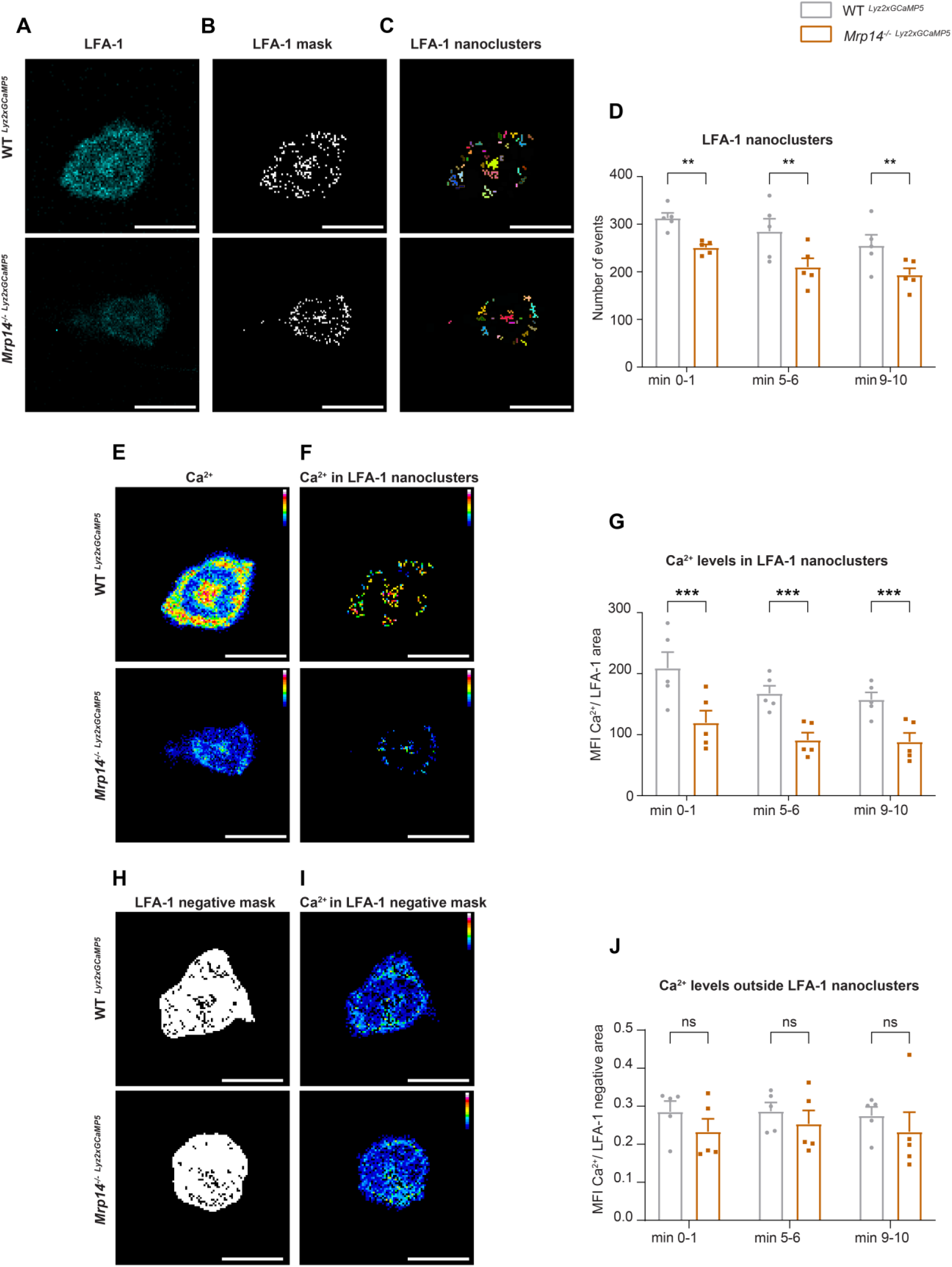
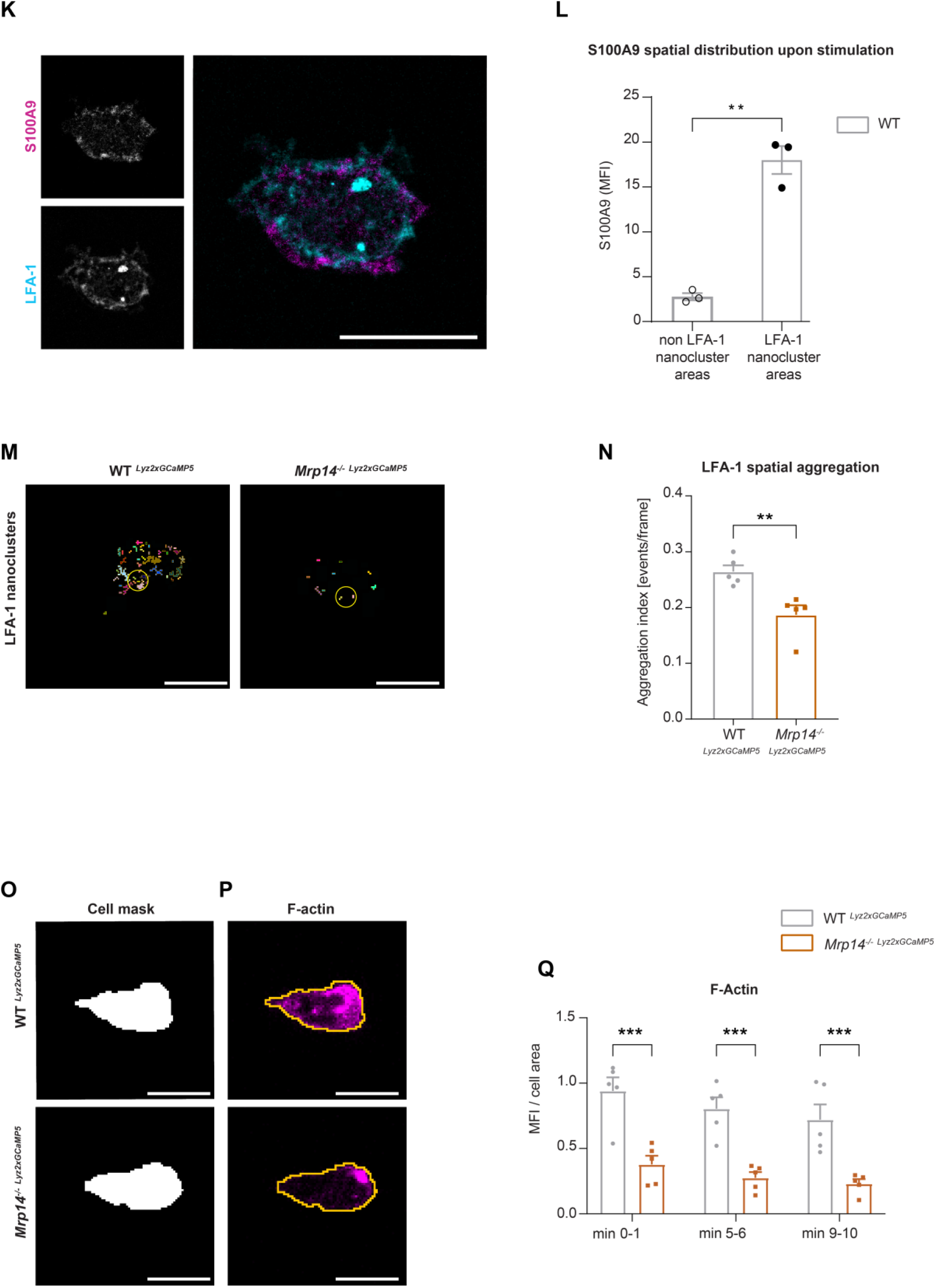
Cytosolic S100A8/A9 drives neutrophil cytoskeletal rearrangement by regulating LFA-1 nanocluster formation and Ca^2+^ availability within the clusters. (A) Representative confocal images of LFA-1 staining in WT *^Lyz2xGCaMP5^*and *Mrp14^−/− Lyz2xGCaMP5^* crawling neutrophils on E-selectin, ICAM-1, and CXCL1 coated flow chambers (scale bar=10μm). (B) Segmentation of LFA-1 signals through automatic thresholding (scale bar=10μm). (C) Size-excluded LFA-1 nanoclusters of 0.15μm^2^ minimum size from previously thresholded images (scale bar=10μm). (D) Single cell analysis of average number of LFA-1 nanoclusters in min 0-1, 5-6 and 9-10 of analysis of WT *^Lyz2xGCaMP5^* and *Mrp14^−/− Lyz2xGCaMP5^*neutrophils [mean+SEM, *n=*5 mice per group, 56 (WT) and 54 (*Mrp14^−/−^*) neutrophils, 2way ANOVA, Sidak’s multiple comparison]. (E) Representative confocal images of Ca^2+^ signals in WT *^Lyz2xGCaMP5^* and *Mrp14^−/− Lyz2xGCaMP5^* neutrophils (scale bar=10μm) and (F) Ca^2+^ signals in the previously segmented LFA-1 nanoclusters (scale bar=10μm). (G) Quantification of subcellular Ca^2+^ levels in the LFA-1 nanocluster area in min 0-1, 5-6 and 9-10 in WT *^Lyz2xGCaMP5^* and *Mrp14*^−/−^ *^Lyz2xGCaMP5^* neutrophils [mean+SEM, *n=*5 mice per group, 56 (WT) and 54 (*Mrp14^−/−^*) cells, 2way ANOVA, Sidak’s multiple comparison]. (H) Segmented LFA-1 cluster negative areas (scale bar=10μm) and (I) representative confocal images of Ca^2+^ signals in the LFA-1 cluster negative areas (scale bar=10μm) [Fig. 4E-I scale bar color code: 0= black, 255=white]. (J) Analysis of cytosolic Ca^2+^ levels in the LFA-1 cluster negative areas in min 0-1, 5-6 and 9-10 of WT *^Lyz2xGCaMP5^* and *Mrp14^−/− Lyz2xGCaMP5^*neutrophils [mean+SEM, *n=*5 mice per group, 56 (WT) and 54 (*Mrp14^−/−^*) neutrophils, 2way ANOVA, Sidak’s multiple comparison]. (K) Representative confocal images showing S100A9 localization at LFA-1 nanocluster areas in stimulated WT neutrophils (scale bar = 10 μm). (L) Quantitative analysis of S100A9 levels in positive LFA-1 nanocluster areas compared to non LFA-1 nanocluster areas in stimulated WT neutrophils. [mean+SEM, *n=*3 mice, 26 (WT) neutrophils, paired Student’s *t*-test]. (M) Representative confocal micrographs of LFA-1 nanocluster spatial aggregation in WT *^Lyz2xGCaMP5^* and *Mrp14^−/− Lyz2xGCaMP5^* neutrophils, within 10μm^2^ area and minimum 10 LFA-1 nanoclusters considered (≥ 10 LFA-1 nanoclusters within 10µm^2^, yellow circles=spatial aggregation area, scale bar=10μm). (N) Analysis of spatially aggregated LFA-1 nanoclusters of WT *^Lyz2xGCaMP5^* and *Mrp14^−/− Lyz2xGCaMP5^* neutrophils [mean+SEM, *n=*5 mice per group, 56 (WT) and 54 (*Mrp14^−/−^*) cells, unpaired Student’s *t*-test]. (O) Segmentation of WT *^Lyz2xGCaMP5^* and *Mrp14^−/− Lyz2xGCaMP5^* neutrophil area through *Lyz2* channel automatic thresholding and (P) representative confocal images of respective F-actin signals. (Q) Analysis of F-actin intensity normalized to the cell area in min 0-1, 5-6 and 9-10 of WT *^Lyz2xGCaMP5^* and *Mrp14^−/− Lyz2xGCaMP5^* neutrophils [mean+SEM, *n=*5 mice per group, 74 (WT) and 66 (*Mrp14^−/−^*) cells, 2way ANOVA, Sidak’s multiple comparison]. ns, not significant; *p≤0.05, **p≤0.01, ***p≤0.001.

In line, overall Ca^2+^ levels under basal conditions (poly-L-lysine coating, static conditions) were similar between WT *^Lyz2xGCaMP5^* and *Mrp14^−/− Lyz2xGCaMP5^* neutrophils (Fig. S3A). Calmodulin levels did not differ between WT *^Lyz2xGCaMP5^* and *Mrp14^−/− Lyz2xGCaMP5^* cells as analyzed by western blot (Fig. S3B).

LFA-1 is known to be rapidly recycled and to spatially redistribute to form a ring like structure that co-clusters with endothelial ICAM-1 during neutrophil migration [33]. To study spatial distribution of LFA-1 nanoclusters (Fig. 4M), we used Ripley’s K function in WT *^Lyz2xGCaMP5^* and *Mrp14^−/− Lyz2xGCaMP5^* neutrophils (Fig. S3C). Ripley’s K is a spatial statistic that compares a given point distribution with a random distribution [34]. WT *^Lyz2xGCaMP5^* neutrophils showed significantly more aggregated LFA-1 nanoclusters within 10μm^2^ area, suitable for LFA-1 enriched pseudopods, compared to *Mrp14^−/− Lyz2xGCaMP5^*neutrophils (Fig. 4N), independent from the total LFA-1 nanocluster number. These results show that in the absence of cytosolic S100A8/A9, LFA-1 nanoclusters are more randomly distributed compared to control and indicate that subcellular redistribution of LFA-1 during migration requires cytosolic S100A8/A9.

Recent work has shown that Ca^2+^ signaling promotes F-actin polymerization at the uropod of polarized neutrophils [13]. Actin waves in turn are known to be important for membrane protrusion formation, neutrophil polarization and firm arrest [35]. Therefore, we examined F-actin dynamics in the presence or absence of cytosolic S100A8/A9. For this, we used the same experimental setting as for the LFA-1 cluster analysis but this time we fluorescently labelled WT *^Lyz2xGCaMP5^* and *Mrp14^−/− Lyz2xGCaMP5^* neutrophils for F-actin. We generated a mask using the myeloid cell marker *Lyz2* (Fig. 4O) and applied the mask to the F-actin channel (Fig. 4P). In line with our previous results on reduced Ca^2+^ levels within LFA-1 adhesion clusters in the absence of S100A8/A9, we found a strongly reduced F-actin signal in *Mrp14^−/− Lyz2xGCaMP5^*neutrophils compared to WT *^Lyz2xGCaMP5^* neutrophils (Fig. 4Q and Movie S4). Total actin levels as determined by western blot analysis did not differ between WT *^Lyz2xGCaMP5^* and *Mrp14^−/− Lyz2xGCaMP5^* neutrophils (Fig. S3D).

Finally, we analyzed the frequency of Ca^2+^ flickers in WT *^Lyz2xGCaMP5^* and *Mrp14^−/− Lyz2xGCaMP5^*neutrophils induced by E-selectin, ICAM-1 and CXCL1 stimulation using high throughput computational analysis. We found an increased number of Ca^2+^ flickers/min in the absence of S100A8/A9 (Fig. S3E and S3F), going along with a shorter duration of the Ca^2+^ event compared to control cells (Fig. S3G and S3H). This finding suggests that cytosolic S100A8/A9 is not only important for local Ca^2+^ supply at focal LFA-1 nanocluster sites, but in addition “stabilizes” Ca^2+^ signaling, preventing fast and uncontrolled Ca^2+^ flickering.

Taken together, these data show that cytosolic S100A8/A9 is indispensable for LFA-1 nanocluster formation and actin-dependent cytoskeletal rearrangements by providing and/or promoting Ca^2+^ supply at the LFA-1 nanocluster sites.

### Cytosolic S100A8/A9 is dispensable for chemokine induced ER store Ca^2+^ release and for the initial phase of SOCE

Our data suggest that intracellular S100A8/A9 is a fundamental regulator of cytosolic Ca^2+^ availability within neutrophils during the recruitment process thereby affecting subcellular LFA-1 and actin dynamics and distribution. Finally, we wanted to study any potential impact of cytosolic S100A8/A9 on Ca^2+^ store release and on Store-Operated Ca^2+^ Entry (SOCE) during neutrophil activation by investigating GPCR induced Ca^2+^ signaling using flow cytometry. First, we investigated Ca^2+^ release from the ER and therefore performed the experiments in absence of extracellular Ca^2+^. We could not detect any differences in CXCL1 induced ER store Ca^2+^ release between WT and *Mrp14^−/−^* cells, indicating that GPCR induced downstream signaling leading to ER store depletion is not affected by the absence of cytosolic S100A8/A9 (Fig. 5A and 5B). In addition, overall basal Ca^2+^ levels (prior to chemokine stimulation) were similar between WT and *Mrp14^−/−^* neutrophils (Fig. 5C).

**Figure 5:**
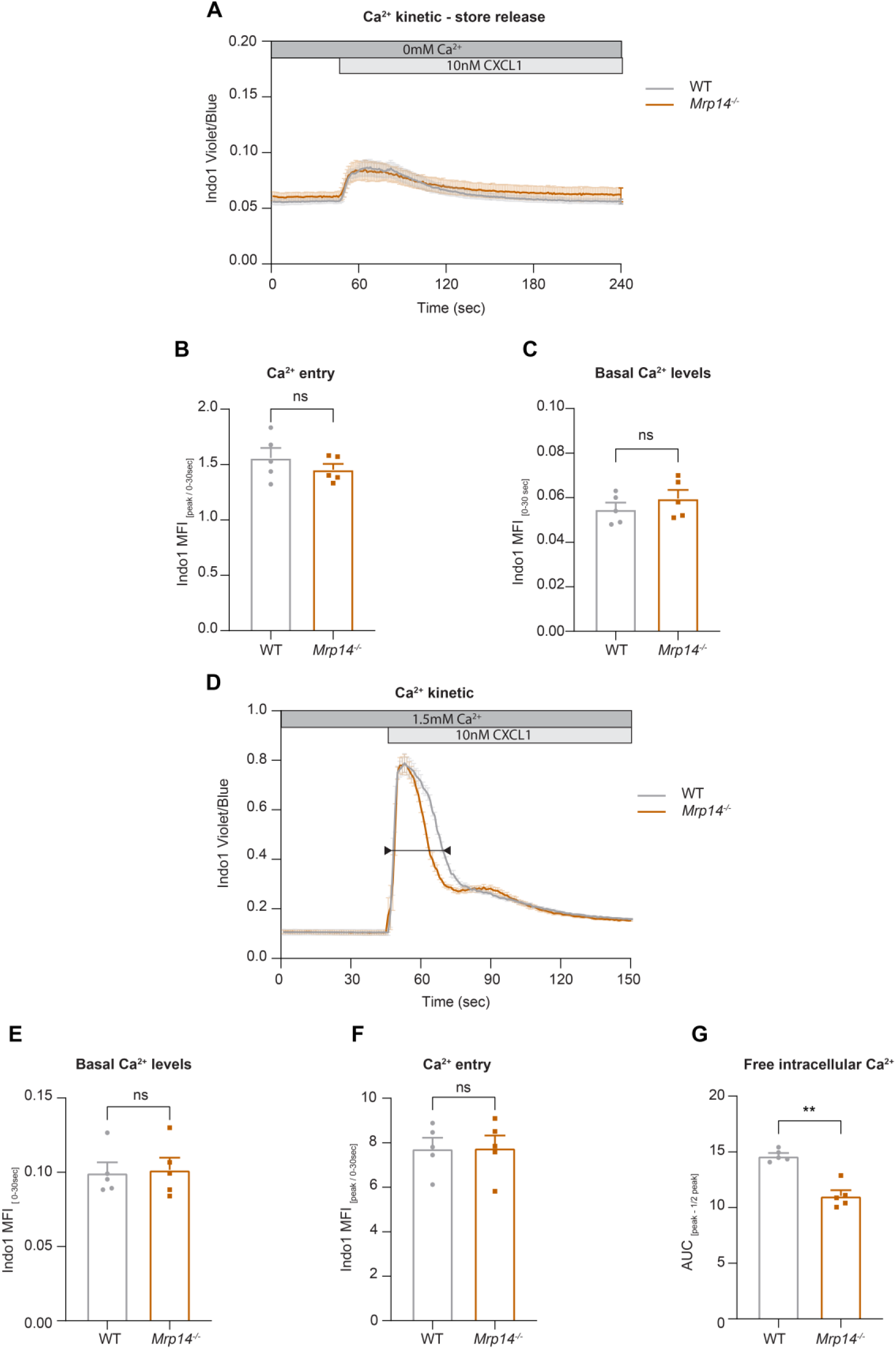
Cytosolic S100A8/A9 is dispensable for chemokine induced ER store Ca^2+^ release and for the initial phase of SOCE. (A) Average flow cytometry kinetic graphs of Ca^2+^ store release in the absence of extracellular Ca^2+^ (Ca^2+^ free medium) in WT and *Mrp14^−/−^* neutrophils upon CXCL1 stimulation (traces are shown as mean+SEM, *n=*5 mice per group). (B) Rapid ER store Ca^2+^ release (MFI _peak_/ MFI _0-30s_) of WT and *Mrp14^−/−^* neutrophils [mean+SEM, *n=*5 mice per group, paired Student’s *t*-test]. (C) Quantification of Ca^2+^ levels under baseline conditions (MFI _0-30s_) [mean+SEM, *n=*5 mice per group, paired Student’s *t*-test]. (D) Average flow cytometry kinetic graphs of Ca^2+^ influx in the presence of extracellular Ca^2+^ (HBSS medium, 1.5mM Ca^2+^) of WT and *Mrp14^−/−^* neutrophils upon CXCL1 stimulation (traces are shown as mean+SEM, *n=*5 mice per group, double-headed arrow represents the time points of quantification). (E) Ca^2+^ levels before CXCL1 stimulation (MFI _0-30s_) [mean+SEM, *n=*5 mice per group, paired Student’s *t*-test]. (F) Quantification of ER store Ca^2+^ release and calcium released activated channel (CRAC) store-operated Ca^2+^ entry (MFI _peak_/ MFI _0-30s_) [mean+SEM, *n=*5 mice per group, paired Student’s *t*-test]. (G) Ca^2+^ influx after CXCL1 stimulation, from peak to peak half-life (AUC _peak – ½ peak_) of WT and *Mrp14^−/−^* neutrophils [mean+SEM, *n=*5 mice per group, paired Student’s *t*-test]. ns, not significant; *p≤0.05, **p≤0.01, ***p≤0.001.

Next, we wanted to investigate whether the absence of cytosolic S100A8/A9 might modify chemokine induced SOCE. Therefore, we stimulated isolated WT and *Mrp14^−/−^* neutrophils with CXCL1 in the presence of extracellular Ca^2+^ (Fig. 5D). Again, basal Ca^2+^ levels were not different between WT and *Mrp14^−/−^* cells (Fig. 5E). Also Ca^2+^ release-activated channels (CRAC) functionality was intact as shown by an identical increase in cytosolic Ca^2+^ amount upon CXCL1 stimulation in WT and *Mrp14^−/−^* neutrophils (Fig. 5F). However, we detected different decay kinetics between WT and *Mrp14^−/−^* neutrophils as *Mrp14^−/−^* neutrophils displayed a steeper decay (Fig. 5G). Taken together, these data suggest that the presence of cytosolic S100A8/A9 is not a prerequisite for chemokine/GPCR induced Ca^2+^ release from ER stores and for the initialization of SOCE via CRAC channels. However, absence of cytosolic S100A8/A9 might disturb Ca^2+^ signaling in a temporal manner.

## DISCUSSION

S100A8/A9 is a Ca^2+^ binding protein, mainly located within the cytosolic compartment of myeloid cells [6, 16]. Once secreted, S100A8/A9 heterodimers exhibit pro-inflammatory effects by engagement with its respective receptors including TLR4 and RAGE on a broad spectrum of effector cells, among them phagocytes, lymphocytes and endothelial cells [16, 20]. In addition, extracellular S100A8/A9 is a well-established biomarker for many acute and chronic inflammatory disorders, including cardiovascular diseases, autoimmune diseases and infections [16, 20, 36]. The tetrameric form of S100A8/A9 was recently shown to have an anti-inflammatory effects during an inflammatory process potentially protecting the organism from overwhelming immune responses [23]. Despite increasing evidence of the pro- and anti-inflammatory effects of secreted S100A8/A9, little is known about its intracellular role in myeloid cells. Here, we show that S100A8/A9 is still abundantly present in the cytosolic compartment of neutrophils even after its active release during inflammation. In addition, we demonstrate that cytosolic S100A8/A9 has a functional impact on neutrophil recruitment during β_2_ integrin outside-in signaling events by ensuring high Ca^2+^ levels at LFA-1 cluster sites independent of its extracellular functions. Neutrophil β_2_ integrin outside-in signaling is known to mediate post-arrest modifications including cytoskeletal rearrangements [22, 37]. *Mrp14^−/−^*cells, which also lack MRP8 in mature cells of the myeloid lineage (functional S100A8/A9 deficient cells) [21, 38], were unable to properly spread, polarize and crawl. This resulted in a marked impairment of adherent neutrophils to withstand physiological shear forces exerted by the circulating blood. Defective outside-in signaling in absence of S100A8/A9 was accompanied by reduced phosphorylation of paxillin and Pyk2, two critical factors involved in the regulation of β_2_ integrin mediated cytoskeletal rearrangements [27]. Of note, *Mrp14^−/−^*myeloid cells have been shown to comprise alterations of cytoskeletal function before [21, 39–43]. In the original publication describing the phenotype of *Mrp14^−/−^*mice, an abnormally polarized cell shape of MRP14 deficient cells was described and therefore a potential role of S100A8/A9 in cytoskeletal reorganization was already hypothesized [21]. In 2014, Vogl et al. demonstrated that cytosolic S100A8/A9 had an impact on the stabilization of microtubules (MTs) via direct interaction of S100A8/A9 with tubulin in resting phagocytes. Upon p38 MAPK and concomitant Ca^2+^ signaling, S100A8/A9 was shown to dissociate from MTs, leading to de-polymerization of MTs thereby allowing neutrophils to transmigrate into inflamed tissue. This might also explain decreased migration of *Mrp14^−/−^* granulocytes in a mouse wound healing model [39]. Additional studies described cytosolic S100A8/A9 to translocate to the membrane and colocalize with vimentin in monocytes upon activation [40], to interact with keratin in epithelial cells [41] and to associate with F-actin localized to lamellipodia in fMLF stimulated neutrophils [42]. Those findings led us to investigate a potential role of cytosolic S100A8/A9 in the Ca^2+^ dependent interplay of plasma membrane located adhesion sites and the cytoskeleton during neutrophil recruitment.

Neutrophil activation during leukocyte recruitment goes along with Ca^2+^ flux initiated e.g. by the engagement of chemokines with G-protein coupled receptors (GPCRs). Subsequently, phospholipase C beta (PLC-β) is activated and leads to the production of Inositol-1,4,5-triphosphate (IP3), which in turn elicits the IP3-receptor in the endoplasmic reticulum (ER), resulting in a rapid Ca^2+^ release from ER stores into the cytoplasm. The decrease in Ca^2+^ concentration in the ER in turn activates the Ca^2+^ sensor stromal interaction molecules (STIM1 and STIM2) triggering the entry of extracellular Ca^2+^ through SOCE mainly via the Ca^2+^-release activated (CRAC) channel ORAI-1 and transient receptor potential (TRP) channels [8, 10, 44]. ORAI-1 is recruited to adhesion cluster sites ensuring high Ca^2+^ levels at the “inflammatory synapse” and rapid rise in intracellular Ca^2+^ concentration, which mediates the assembly of cytoskeletal adaptor proteins to integrin tails and allows the onset of pseudopod formation [14, 45]. The importance of localized Ca^2+^ availability in subcellular domains has also been shown in T-cells during the engagement with antigen presenting cells within the immunological synapse. In T-cells, mitochondria play a central role as Ca^2+^ buffers and as Ca^2+^ conductors that collect cytosolic Ca^2+^ at the entry site, (i.e. through open CRAC channels located at the immunological synapse) and distribute it throughout the cytosol. [46]. Here we show that in neutrophils cytosolic S100A8/A9 colocalizes with LFA-1 during intravascular adhesion and increases and stabilizes Ca^2+^ availability at the LFA-1 nanocluster sites, mediating spatial clustering of LFA-1 and sustained polymerization of F-actin, both essential steps for efficient neutrophil adhesion strengthening. In addition, presence of cytosolic S100A8/A9 stabilizes duration of Ca^2+^ signals within the cells, as WT cells displayed longer frequencies of Ca^2+^ events with less flickers min^-1^, which might in addition be important for the stability of the inflammatory synapse.

As reported earlier, chemokine induced Ca^2+^ influx through SOCE at the plasma membrane is indispensable for the activation of high affinity β_2_ integrins [11]. This early step during leukocyte recruitment (inside-out signaling) was not affected in absence of cytosolic S100A8/A9. In line, *Mrp14^−/−^* cells displayed similar CXCL1 induced Ca^2+^ fluxes compared to WT cells. Ca^2+^ release from intracellular stores and initial phases of SOCE were fully functional, as shown by flow cytometry of Indo-1 dye loaded neutrophils. These findings are in accordance with a study by Hobbs et al., which also described normal Ca^2+^ influx in S100A8/A9 deficient neutrophils induced by the chemokine MIP-2 [47]. However, we found an impact of cytosolic S100A8/A9 in sustaining high Ca^2+^ concentrations, as Ca^2+^ fluxes decreased faster in the absence of S100A8/A9. Whether this faster decrease is mediated through a direct effect of cytosolic S100A8/A9 on SOCE or through a potential buffer capacity of cytosolic S100A8/A9 needs to be further investigated. Hobbs et al. proposed no impact of S100A9 deletion in the recruitment of neutrophils by using a thioglycolate induced peritonitis model. However, peritoneal neutrophil emigration was shown to be rather independent of LFA-1 [48, 49], whereas extravasation into cremaster muscle tissue strongly relies on the β_2_ integrin LFA-1 and integrin clustering [50].

Taken together, we identified a critical role of cytosolic S100A8/A9 in neutrophil recruitment. We show that its absence leads to reduced Ca^2+^ signaling and impaired sustained Ca^2+^ supply at LFA-1 nanocluster sites. Attenuated Ca^2+^ signatures in turn affect β_2_ integrin dependent cytoskeletal rearrangements and substantially compromises neutrophil recruitment during the inflammatory response. These findings uncover cytosolic S100A8/A9 as a potentially interesting therapeutic target to reduce neutrophil recruitment during inflammatory disorders with unwanted overwhelming neutrophil influx.

## MATERIALS AND METHODS

### Mice

C57BL/6 wildtype (WT) mice were purchased from Charles Rivers Laboratories (Sulzfeld, Germany). *Mrp14^−/−^* (functional double S100A8 and S100A9 ko animals) mice were kindly provided by Johannes Roth (Institute for Immunology, Muenster, Germany). *B6;129S6-Polr2atm1(CAG-GCaMP5g-tdTomato)* crossbred with Lyz2^Cre^ (*GCaMP5xWT*) were kindly provided by Konstantin Stark (LMU, Munich, Germany) and crossbred with *Mrp14^−/−^* mice (GCaMP5x*Mrp14^−/−^*). All mice were housed at the Biomedical Center, LMU Munich, Planegg-Martinsried, Germany. Male and/or female mice (8–25 weeks old) were used for all experiments. Animal experiments were approved by the Regierung von Oberbayern (AZ.: ROB-55.2-2532.Vet_02-17-102 and ROB-55.2-2532.Vet_02-18-22) and carried out in accordance with the guidelines from Directive 2010/63/EU. For in vivo experiments, mice were anaesthetized via i.p. injection using a combination of ketamine/xylazine (125mg kg^-1^ and 12.5mg kg^-1^ body weight, respectively in a volume of 0.1mL NaCl per 8g body weight). All mice were sacrificed at the end of the experiment by cervical dislocation.

### Neutrophil isolation

Bone marrow neutrophils were isolated using the EasySep Mouse Neutrophil Enrichment Kit according to the manufacturer’s instructions (STEMCELL Technologies). Isolated neutrophils were then resuspended in HBSS buffer [containing 0.1% of glucose, 1mM CaCl_2_, 1mM MgCl_2_, 0.25% BSA, and 10mM HEPES (Sigma-Aldrich), pH7.4, complete HBSS].

### S100A8/A9 ELISA

In vitro release of S100A8/A9 was performed as described before [51]. Briefly, bone marrow neutrophils were isolated from WT mice. 24 well-plates were coated with recombinant murine (rm) E-selectin (rmCD62E-Fc chimera, 10µg mL^-1^, R&D Systems) or PBS/0.1% BSA at 4°C overnight, blocked with PBS/5% casein (Sigma-Aldrich) and washed twice with PBS. 5×10^5^ neutrophils were reconstituted in complete HBSS buffer and incubated under shaking conditions on the coated slides for 10min at 37°C and 5% CO2. To assess the total intracellular S100A8/A9 levels, cells were lysed in 2% Triton X-100 (Applichem). Finally, cellular supernatants were analyzed by Enzyme-Linked Immunosorbent Assay (ELISA) to determine the concentrations of S100A8/A9.

### Murine cremaster muscle models

Leukocyte recruitment was investigated by intravital microscopy in inflamed cremaster muscle venules as reported previously [52]. Shortly, intrascrotal (i.s.) injection of rmTNF-α (500ng, R&D Systems) was applied to WT and *Mrp14^−/−^* mice in order to induce an acute inflammation in the cremaster muscle. Two hours after injection, the carotid artery of anaesthetized mice was catheterized for later blood sampling (ProCyte Dx; IDEXX Laboratories) or intra-arterial (i.a.) injection. Thereafter, the cremaster muscle was exteriorized and intravital microscopy was conducted on an OlympusBX51 WI microscope, equipped with a 40x objective (Olympus, 0.8NA, water immersion objective) and a CCD camera (KAPPA CF 8 HS). Post-capillary venules were recorded using VirtualDub software for later analysis. Rolling flux fraction, number of adherent cells mm^−2^, vessel diameter and vessel length were analyzed using FIJI software [53]. During the entire experiment, the cremaster muscle was superfused with thermo-controlled bicarbonate buffer as described earlier [54]. Centerline blood flow velocity in each venule was measured with a dual photodiode (Circusoft Instrumentation). Subsequently, cremaster muscles were removed, fixed in 4% PFA solution O.N. at 4°C and the next day stained with Giemsa (Merck) to assess the number of perivascular neutrophils. The tissues were mounted in Eukytt mounting medium and covered with a 170μm coverslip. Neutrophils were discriminated from other leukocyte subpopulations based on nuclear shape and granularity of the cytosol. The analysis of transmigrated leukocytes was carried out at the Core Facility Bioimaging of the Biomedical Center with a Leica DM2500 transmission bright field microscope, equipped with a 100x, 1.4 NA, oil immersion objective and a Leica DMC2900 CMOS camera. Resulting images had 2048×1536 pixels and a pixel size of 58nm.

For rescue experiments, we adopted either the TNF-α-induced inflammation model as described above or the trauma-induced inflammation model of the mouse cremaster muscle. In the trauma model, sterile inflammation was induced by opening and exteriorizing the cremaster muscle without application of any stimulus. Intravital microscopy was conducted as described above. After finding an appropriate spot, the same vessel was recorded before and after injection of mutant murine S100A8/S100A9N70AE79A (S100A8/A9^mut^, aa exchange N70A and E79A, 50µg mouse^-1^ in 100µL, provided by Thomas Vogl, University of Muenster, Germany) and the number of adherent cells mm^-2^ were counted pre and post injection in WT and *Mrp14^−/−^*mice.

### S100A8/A9 intracellular staining

For the analysis of cytosolic S100A8/A9 levels, TNF-α stimulation of the mouse cremaster muscle was carried out as described above. Subsequently, cremaster muscles were removed, fixed in 4% PFA solution, and immunofluorescence staining for PECAM-1 (AlexaFluor488 labelled primary monoclonal rat antibody, 5μg mL^-1^, MEC13.3, BioLegend) and S100A9 (Cy5.5 directly labelled, 5μg mL^-1^, clone 322, provided by Thomas Vogl) was conducted. Stained samples were mounted in Vectashield mounting medium, covered with a 0.17μm coverslip and imaged by confocal microscopy at the Core Facility Bioimaging of the Biomedical Center, LMU Munich, with an upright Leica SP8X WLL microscope, equipped with a HC PL APO 40x/1.30 NA oil immersion objective. AF488 was excited with 488nm, Cy5.5 with 543nm. Detection windows were 500 – 568 and 550 – 640 nm, respectively. Both channels were recorded sequentially. Hybrid photodetectors were used to record images with 512×512 pixels with a pixel size of 0.427μm. Single cell analysis was carried out by FIJI software using macros as follows: MAX projection of Z-stacks were created and neutrophils were segmented by thresholding using S100A8/A9 signal. Then, cell masks were applied back to the original images and S100A8/A9 mean fluorescence intensity (MFI) averaged on stack slices. Finally, S100A8/A9 MFIs were analyzed from intravascular and extravasated neutrophils.

### Neutrophil surface marker staining

Peripheral blood from WT and *Mrp14^−/−^* mice was harvested and erythrocytes were lysed with lysing solution (BD FACS™). Samples were stained for CD18-FITC (5μg mL^-1^; C71/16; Pharmigen), CD11a-APC (2μg mL^-1^; M17/4; eBioscience), CD11b-BV510 (0.3μg mL^-1^; M1/70; BioLegend), CD62L-FITC (5μg mL^-1^; MEL-14; BioLegend), PSGL1-PE (2μg mL^-1^; 2PH1; Pharmigen), CXCR2-APC (5μg mL^-1^; 242216; R&D Systems), CD44-BV570 (0.3μg mL^-1^; IM7; BioLegend). Respective isotype controls were used: IgG2a-FITC (5μg mL^-1^; RTK2759; BioLegend), IgG2a-APC (2μg mL^-1^; RTK2758; BioLegend), IgG2b-BV510 (0.3μg mL^-1^; RTK4530; BioLegend), IgG1-PE (2μg mL^-1^; eBRG1; eBioscience), IgG2b-BV570 (0.3μg mL^-1^; RTK4530; BioLegend). Neutrophils were defined as Ly6G^+^ cells (0.8μg mL^-1^; 1A8; BioLegend).

### Neutrophil adhesion ex vivo

Flow chamber assays were carried out as previously described [22]. Briefly, rectangular borosilicate glass capillaries (0.04×0.4mm; VitroCom) were coated with a combination of rmE-selectin (CD62E Fc chimera; 20µg mL^−1^; R&D Systems), rmICAM-1 (ICAM-1 Fc chimera; 15µg mL^−1^; R&D Systems), and rmCXCL1 (15µg mL^−1^; Peprotech) for 3h at RT and blocked with PBS/5% casein (Sigma-Aldrich) over night at 4°C. WT and *Mrp14^−/−^* whole blood was perfused through the microflow chamber via a carotid artery catheter of anesthetized mice at varying shear stress levels. Movies were recorded on an OlympusBX51 WI microscope with a 20x, 0.95NA, water immersion objective and a CCD camera (KAPPA CF 8 HS) with VirtualDub software [55]. Resulting images had 768×576 pixels and a pixel size of 0.33μm .Number of rolling and adherent leukocytes/field of view (FOV) were counted using Fiji software, over one-minute time window after 6min of blood infusion.

### β_2_ integrin activation assay

β_2_ integrin activation was determined through a modified soluble ICAM-1 binding assay [6]. Bone marrow murine neutrophils were isolated as described above. Enriched neutrophils (1.5×10^6^) were incubated and stained with rmICAM-1 Fc chimera (40µg mL^−1^, R&D Systems), IgG-Fc-biotin (12.5µg mL^−1^; eBioscience), and streptavidin-PerCP-Cy5.5 (2µg mL^−1^; BioLegend). Then, cells were stimulated with rmCXCL1 (10nM) or PBS (control) in complete HBSS buffer for 5min at 37°C. The amount of bound rmICAM-1 to the β_2_ integrin was assessed by flow cytometry (CytoFlex S, Beckmann Coulter) and the median shift relative to the control was analyzed by FlowJo software.

### Static adhesion assay

Neutrophil static adhesion assay was performed as previously described [56]. Shortly, 96-well plates were coated with rmICAM-1 (3µg mL^-1^) over night at 4°C and washed with PBS. Neutrophils were resuspended in complete HBSS and seeded at 1×10^5^ cells per well. Cells were allowed to settle for 5min at 37°C and stimulated with 10nM rmCXCL1 or PBS (control) for 10min at 37°C. Using a standard curve, adherent neutrophils were calculated as percentage of total cells added. Standard curve preparation was done by adding 100%, 80%, 60%, 40%, 20%, and 10% of the cell suspension on poly-L-lysine coated wells (100µg mL^-1^) in triplicates. Non adherent cells were washed away while adherent cells were fixed with 1% glutaraldehyde and stained with 0.1% crystal violet solution (Sigma-Aldrich). Absorption at 590nm was measured with a microplate reader (PowerWave HT, Biotek, USA) after lysis of cells with 10% acetic acid solution, as previously described [57].

### Spreading assay

To study neutrophil spreading, rectangular borosilicate glass capillaries (0.04×0.40mm; VitroCom) were coated with rmE-selectin (CD62E Fc chimera; 20µg mL^−1^), rmICAM-1 (15µg mL^−1^), and rmCXCL1 (15µg mL^−1^) for 3 hours at RT and blocked with PBS/5% casein over night at 4°C. Bone marrow neutrophils were matured in RPMI 1640 (Sigma-Aldrich) containing FCS (10%, Sigma-Aldrich), GlutaMAX (1%, ThermoFisher), Penicillin-Streptomycin solution (1%, Corning®) and supplemented with 20% WEHI-3B-conditioned medium over night at 37°C and applied into the flow chamber at a shear stress level of 1dyne cm^−2^ using a high-precision syringe pump (Harvard Apparatus, Holliston, Massachusetts, USA). Cells were incubated with Fc-block (murine TruStain FcX; BioLegend) for 5min at RT before being introduced into the chambers. Spreading behavior of the cells was observed and recorded on a Zeiss Axioskop2 with a 20x, 0.5NA water immersion objectibve and a Hitachi KP-M1AP camera with VirtualDub. Resulting images had 1360×1024 pixels and a pixel size of 600nm. Cell shape changes were quantified using FIJI software, analyzing cell area, perimeter, circularity 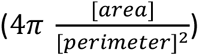 and solidity 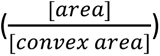.

### Crawling assay

15µ-Slides VI^0.1^ (Ibidi) were coated with a combination of rmE-selectin (20µg mL^−1^), rmICAM-1 (15µg mL^−1^), and rmCXCL1 (15µg mL^−1^) for 3h at RT and blocked with PBS/5% casein over night at 4°C. Overnight matured bone marrow neutrophils from WT and *Mrp14^−/−^* mice were resuspended in complete HBSS at 1×10^6^ mL^-1^, introduced into the chambers and allowed to settle and adhere for 3min until flow was applied (2dyne cm^-2^) using a high-precision perfusion pump. Experiments were conducted on a ZEISS, AXIOVERT 200 microscope, provided with a ZEISS 20x objective [0.25NA], and a SPOT RT ST Camera. MetaMorph software was used to generate time-lapse movies for later analysis. 20min of neutrophil crawling under flow were analyzed using FIJI software [53] and chemotaxis tool plugin (Ibidi).

### Paxillin and Pyk2 phosphorylation

Paxillin and Pyk2 phosphorylation was investigated as previously described [22]. Briefly, 2×10^6^ WT or *Mrp14^−/−^*bone marrow murine neutrophils were seeded on rmICAM-1 coated wells (15µg mL^−1^) for 5min and stimulated with rmCXCL1 (10nM) for 5min at 37°C. Cells were then lysed with lysis buffer [containing 150mM NaCl, 1% Triton X-100, 0.5% Sodium deoxycholate (Sigma-Aldrich), 50mM Tris–HCl pH7.3 (Merck), 2mM EDTA (Merck) supplemented with protease (Roche), phosphatase inhibitors (Sigma-Aldrich) and 1xLaemmli sample buffer] and boiled (95°C, 5min). Cell lysates were resolved by SDS–PAGE and electrophoretically transferred onto PVDF membranes. After subsequent blocking (LI-COR blocking solution), membranes were incubated with the following antibodies for later detection and analysis using the Odyssey® CLx Imaging System and Image Studio software: rabbit α-mouse phospho-Paxillin (Tyr118) or rabbit α-mouse Paxillin and rabbit α-mouse phospho-Pyk2 (Tyr402) or rabbit α-mouse Pyk2 (all Cell Signaling). Goat-α-rabbit IRDye 800RD was used as secondary antibody (Licor).

### Detachment assays

To investigate shear resistance, rectangular borosilicate glass capillaries (0.04×0.40mm; VitroCom) were coated with rmE-selectin (CD62E Fc chimera; 20µg mL^−1^), rmICAM-1 (15µg mL^−1^), and rmCXCL1 (15µg mL^−1^) for 3 hours at RT and blocked with 5% casein over night at 4°C. Whole blood from WT and *Mrp14^−/−^* mice was perfused in the coated flow chambers via the cannulated carotid artery, where neutrophils were allowed to attach for 3min. Then, flow was applied through a high-precision perfusion pump and detachment assays performed over 10min with increasing shear stress (34 – 272 dyne cm^-2^) every 30s. Experiments were recorded by time-lapse movies using the upright Zeiss Axioskop2 with the 20x, 0.5 NA water immersion objective as described above. Number of attached cells was counted at the end of each step.

### LFA-1 clustering, S100A8/A9 distribution, Ca^2+^ localization and F-actin signature during neutrophil crawling under flow

15µ-Slides VI^0.1^ (Ibidi) were used to study LFA-1 clustering, Ca^2+^ localization and F-actin signature during neutrophil crawling. Flow chambers were coated and blocked as described above. 2×10^6^ isolated neutrophils from WT *^Lyz2xGCaMP5^* or *Mrp14^-/-Lyz2xGCaMP5^* were stained with in-house AlexaFluor647 labelled (Antibody Labeling Kit, Invitrogen™) monoclonal anti LFA-1 rat antibody (5μg mL^-1^, 2D7, BD Pharmingen) for 10min prior to the experiment or SiR-actin (200nM, Spirochrome™) O.N., respectively. Cells were seeded in the chambers and allowed to settle for 2min before flow was applied (2dyne cm^-2^) using a high-precision perfusion pump. Samples were imaged by confocal microscopy at the core facility Bioimaging of the Biomedical Center with an inverted Leica SP8X WLL microscope, equipped with a HC PL APO 40x/1.30 NA oil immersion objective. Observation was at 37°C. Hybrid photodetectors were used to record images with 512×512 pixels and a pixel size of 0.284μm. GCaMP5-GFP was excited with 488nm, AF647 or SiR-Actin with 633nm. Detection windows were 498 – 540 and 649 – 710nm, respectively. For movies, one image was recorded every 0.44 seconds or every 2 seconds, over 10min. Automated single cell analysis was performed using macros with Fiji software, for minute 0-1, minute 5-6 and minute 9-10 of each recording. For the LFA-1 nanocluster analysis, the LFA-1 channel was automatically segmented and ROIs of a minimum size of 0.15μm^2^ were considered as LFA-1 nanosclusters, as reported earlier [32]. This represented a minimum size of 2 pixels in our analysis. The number of clusters was averaged for each analyzed time point (min 0-1, min 5-6 and min 9-10). For the subcellular Ca^2+^ analysis at the LFA-1 cluster sites, the LFA-1 segmented channel was applied to the Ca^2+^ channel and Ca^2+^ events in the selected ROIs were determined, normalized to the LFA-1 areas, and averaged over each minute of analysis. For the Ca^2+^ analysis in the negative LFA-1 area, we again adopted semi-automated single cell analysis and subtracted the LFA-1 mask from the *Lyz2* mask in order to obtain “LFA-1 cluster negative masks”. Later, the “LFA-1 cluster negative masks” were applied to the Ca^2+^ channel and Ca^2+^ intensities were measured, normalized to the “LFA-1 cluster negative masks” and averaged over each minute of analysis. For the analysis of S100A9 distribution at LFA-1 nanocluster areas, WT neutrophils were stained with CellTracker Green CMFDA (10µM, Invitrogen™) for 45min and in-house AlexaFluor647-labeled monoclonal anti-LFA-1 rat antibody (5μg/mL, 2D7, BD Pharmingen) for 10min prior to the experiment. The cells were then seeded in chambers and allowed to settle for 3min, before applying continuous flow (2 dyne cm^−^²) using a high-precision perfusion pump for 10min. After the flow, the cells were fixed, permeabilized, and stained overnight at 4°C for intracellular S100A9, followed by counterstaining with DAPI. A semi-automated single-cell analysis was performed to measure S100A9 intensity in the LFA-1 nanocluster areas (obtained as described above) and in the negative LFA-1 nanocluster areas (determined using the same procedure but with CellTracker Green as the cell mask).

For the F-actin analysis, the *Lyz2* channel was automatically segmented to obtain a cell mask and applied to the F-actin channel. F-actin intensities were measured and averaged over each minute of analysis as described above.

### Ca^2+^ store release and Ca^2+^ influx measurement – flow cytometry

Ca^2+^ store release and Ca^2+^ influx was analyzed by flow cytometry through an adapted protocol [58]. WT and *Mrp14^−/−^* bone marrow neutrophils (2.5×10^6^ mL^-1^) were resuspended in PBS and loaded with 3 µM Indo-1 AM (Invitrogen™) for 45min at 37°C. Cells were washed, resuspended in complete HBSS buffer (2.5×10^6^ mL^-1^) and stained with an anti Ly6G-APC antibody (1μg mL^− 1^, 1A8, BioLegend) and with the Fixable Viability Dye eFluor™ 780 (1:1000; eBioscience™). Cells (2×10^5^) were incubated for 2min at 37°C and 10nM CXCL1 was placed on the side of the FACS tube in a 2μL droplet form. The cells were analyzed at the flow cytometry core facility of the biomedical center with a BD LSRFortessa^TM^ flow cytometer. Samples were recorded for 45 seconds to establish a baseline. Afterwards, CXCL1 stimulation was initiated by tapping the tube with subsequent fall of the drop into the cell suspension while continuously recording Indo-1 AM signals from neutrophils over time. Data were analyzed using FlowJo software. Calcium levels are expressed as relative ratios of fluorescence emission at 375nm/525nm (calcium bound/calcium unbound) and Ca^2+^ signatures quantified as AUC of kinetic averages. To measure Ca^2+^ store release only, Ca^2+^ free medium was used.

### Spatial distribution analysis of LFA-1 nanoclusters

To evaluate the spatial distribution of LFA-1 nanoclusters in neutrophils, Ripley’s K statistics [34] was calculated for every time point in every experiment with radii between 0.5μm and 5.5μm. For every radius r, we calculated the K(r) value as follows:

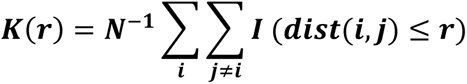

Where *i* and *j* are two different LFA-1 nanocluster locations, *I* is the indicator function which is 1 if the content within the parentheses is “True” and 0 if the content is “False”, and N is a normalization constant. For ‘dist’, the Euclidian distance was chosen and calculated via the “pairwise_distances” from sklearn [59]. The sampling part of Ripley’s K statistic was done by drawing random locations as LFA-1 nanocluster events from the cell surface. To make Ripley’s K results comparable between different experiments, we normalized K(r) values such that the random sampling upper bound, calculated for every experiment, was set to 1, and the random sampling lower bound was set to −1. Thus, every normalized value between −1 and 1 is within random borders, i.e. not distinguishable from a random spatial distribution. Values above 1 indicate aggregated LFA-1 nanoclusters and values below −1 indicate dispersed LFA-1 nanoclusters. At least 10 LFA-1 nanoclusters were considered for the spatial aggregation analysis. The number of identified aggregated LFA-1 nanoclusters (values above 1) was counted and averaged for every condition, resulting in an aggregation index.

### Frequency and duration of Ca^2+^ oscillations

After the recording, Ca^2+^ mean intensities of the cells were calculated over time, counting each cell as an individual ROI. Data was imported into a previously described custom analysis pipeline for Ca^2+^ imaging data [60]. Briefly, the Ca^2+^ mean intensities were sequentially filtered according to the standard values of the pipeline, considering only events with a z-score of at least 3 (p<0.01). From those events, a graph was constructed to detect superimposed events. Properties of the events, AUC or half-width, were used in the calculations afterwards. The code of the analysis pipeline can be accessed in the corresponding repository at https://github.com/szarma/Physio_Ca.

### Calmodulin and β-Actin Western Blotting

WT and *Mrp14^−/−^* bone marrow murine neutrophils (1×10^6^) were isolated as described above and lysed with lysis buffer and boiled (95°C, 5min). Cell lysates were resolved by SDS–PAGE and electrophoretically transferred onto PVDF membranes. After subsequent blocking (LI-COR blocking solution), membranes were incubated with the following antibodies for later detection and analysis using the Odyssey® CLx Imaging System and Image Studio software. Rabbit α-mouse Calmodulin (5μg mL^-1^, CellSignaling), rabbit α-mouse β-actin (1μg mL^-1^, CellSignaling) and mouse α-mouse GAPDH (1μg mL^-1^, Merck/Millipore), goat-α-mouse IRDye 680RD, and goat-α-rabbit IRDye800CW-coupled secondary antibodies (1μg mL^-1^, LI-COR).

### Statistics

Data are presented as mean +SEM, as cumulative distribution or representative images, as depicted in the figure legends. Group sizes were selected based on experimental setup. Data were analyzed and illustrated using GraphPad Prism 9 software. Statistical tests were performed according to the number of groups being compared. For pairwise comparison of experimental groups, a paired/unpaired Student’s t-test and for more than two groups, a one-way or two-way analysis of variance (ANOVA) with either Tukey’s (one-way ANOVA) or Sidak’s (two-way ANOVA) *post-hoc* test with repeated measurements were performed, respectively. p-values <0.05 were considered statistically significant and indicated as follows: ∗p<0.05; ∗∗p<0.01; ∗∗∗p<0.001.

## Supporting information

Supplementary Files (figures with legends and tables)

## Acknowledgements

We thank Dorothee Gössel, Anke Lübeck, Sabine D’Avis, Susanne Bierschenk and Jennifer Troung for excellent technical assistance, and the core facility Flow Cytometry at the Biomedical Center, LMU, Planegg-Martinsried, Germany.

## Funding

This work was supported by the German Research Foundation (DFG) collaborative research grants: SFB914, projects A02 (B.W.), B01 (M.S.), B11 (M.P.), TRR-332 projects C2 (M.S.), C3 (B.W.), C7 (T.V.), B5 (J.R.) and the TRR-359 project B2 (M.S. and R.I.).

## Author contributions

M.N. designed and conducted experiments, analyzed and interpreted data, and wrote the manuscript. R.I. and I.R. designed experiments, acquired and analyzed data V.L. performed the integrin spatial cluster analysis. J.P. performed the calcium frequency distribution analysis. M.Sa and A.Y. analyzed results. T.V. provided critical reagents (S100A8/A9^mut^, anti-S100A9-Cy5.5) and analyzed ELISA samples. J.R., M.S.R, C.M., and B.W. provided their expertise and conceptual advice. M.S. designed experiments and wrote the manuscript. M.P. designed experiments, wrote the manuscript and supervised the work. All authors discussed the results, commented on and approved the manuscript.

## Competing interests

The authors declare no competing interests.

## Data and materials availability

All data and materials that support the findings of this study are available from the corresponding author upon request. Custom codes developed for data analysis and visualization are available at https://github.com/Napo93/AG-Sperandio-MACROS, https://github.com/marrlab/Spatial_CA#spatial_ca and https://github.com/szarma/Physio_Ca. Software and parameters used are stated in the Methods with further details.

## Notes

https://jmp.sh/s/0Eepmup2xJ3WOTpCPzWX

https://jmp.sh/kgXT7AUx

https://jmp.sh/pOQyv0Rg

https://jmp.sh/XxIp8ofF

### Competing Interest Statement

The authors have declared no competing interest.

### Summary of Updates

New text added for revision in red (Result, Discussion, Figure Legend and Method sections); Fig.3I revised and updated with more data and changed representative images (western blot); Fig.4 revised and result section on Fig.4 updated to add colocalization analysis between S100A8/A9 and LFA-1 nanoclusters (Fig.4K and 4L new). Method section, figure legend and discussion for Fig.4K and 4L updated; Fig.5A and 5D updated to add colored line legend; Supplementary Fig. 3e and 3G updated to add colored line legend; Gene locus name (S100a8/a9) changed to lowercase letters in abstract; Supplemental files updated and Supplementary Videos added; Reference list updated (nr.#19 is now published)

