## Supplementary Files (figures with legends and tables) for "Cytosolic S100A8/A9 promotes Ca^2+^ supply at LFA-1 adhesion clusters during neutrophil recruitment"

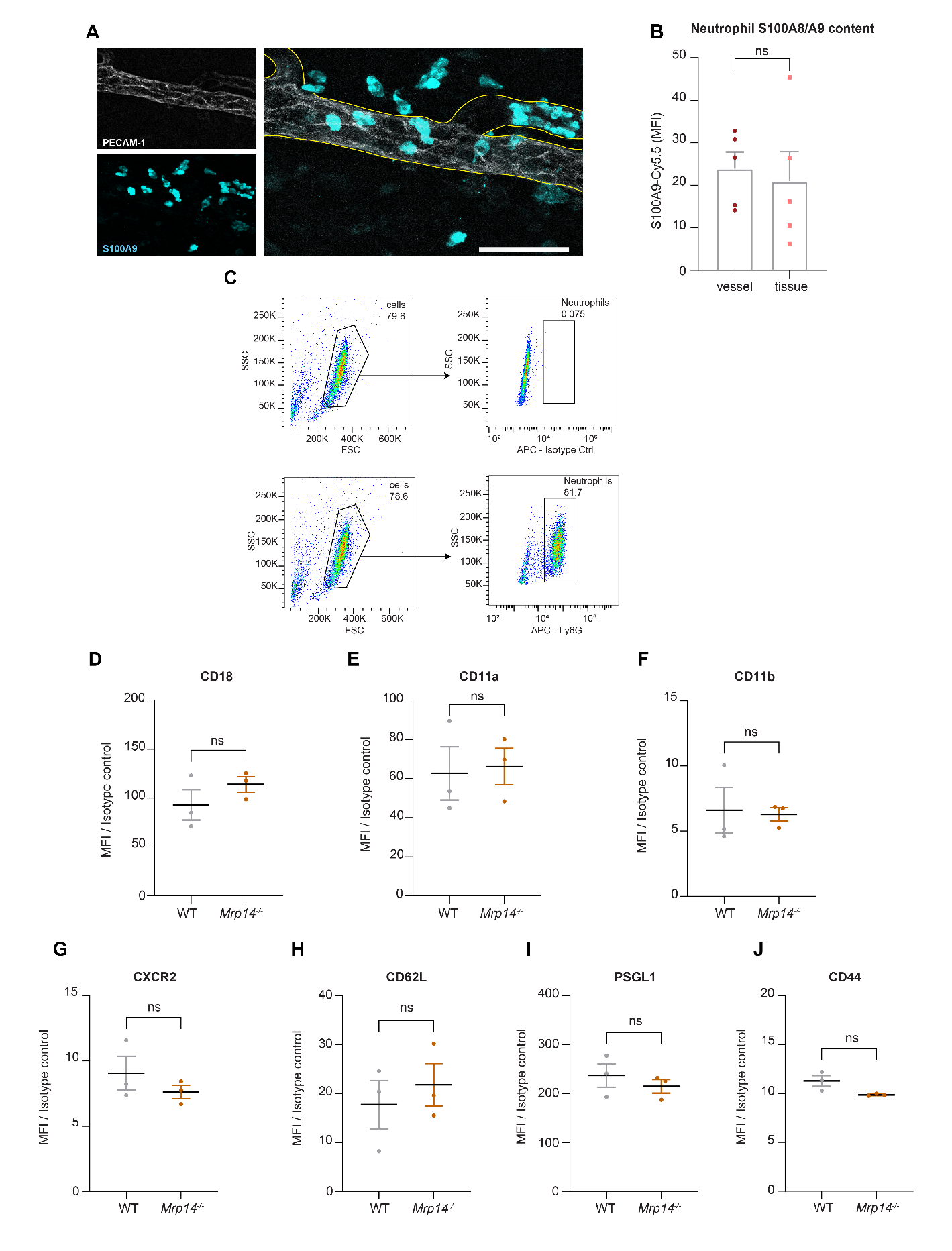
**SUPPLEMENTARY FIGURE 1**

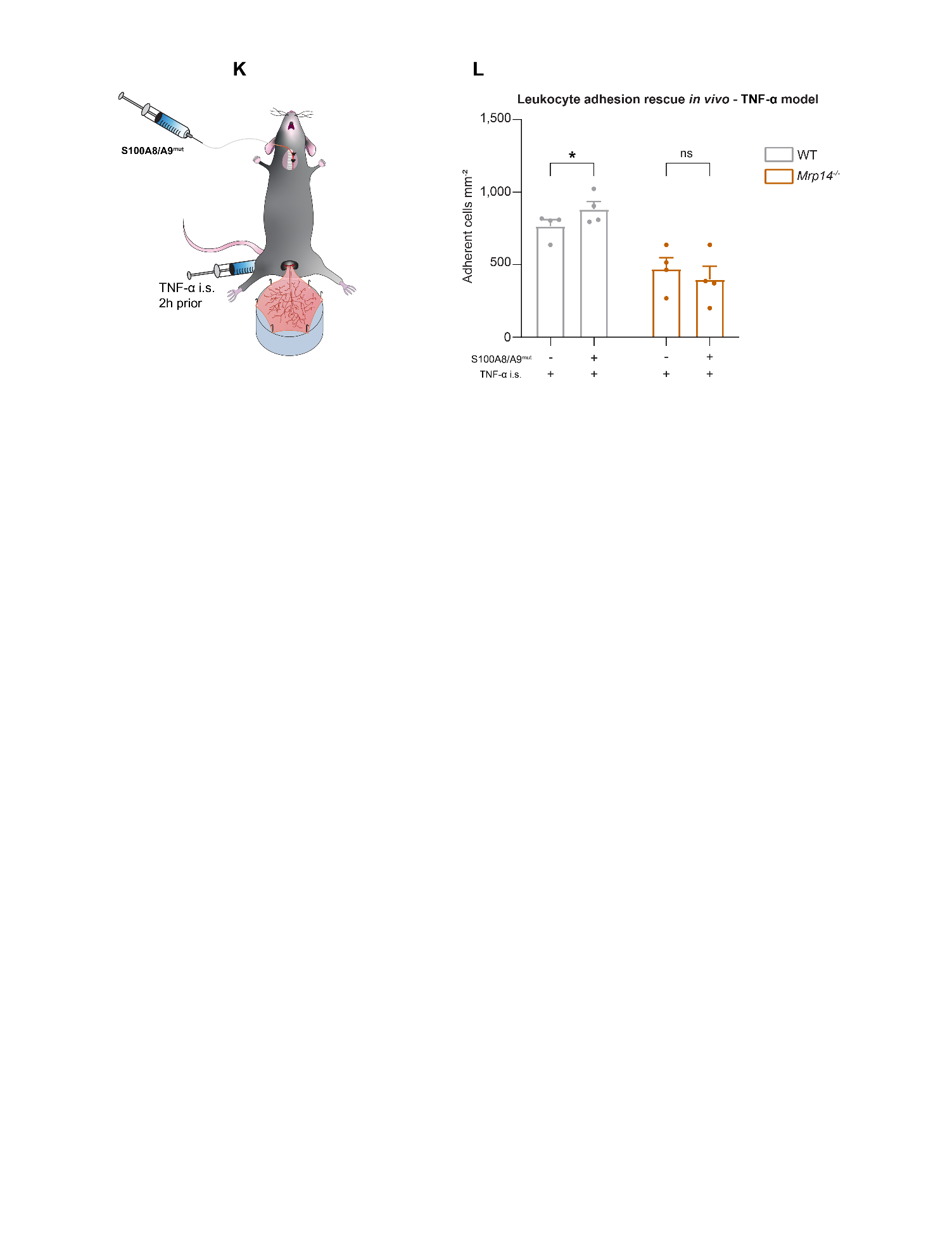

**Figure S1: Cytosolic S100A8/A9 is indispensable for neutrophil recruitment in vivo.** (A) Representative confocal images of S100A9 intensity in intravascular and extravascular neutrophils after TNF-α stimulation of WT cremaster muscle tissues (scale bar=50μm) and (B) quantification [mean+SEM, *n=*5 mice per group, 602 (intravascular) and 326 (extravasated) neutrophils, unpaired Student’s *t*-test]. (C) WT neutrophils’ purity assessment and gating strategy (equivalent for *Mrp14^-/-^* neutrophils) and flow cytometry analysis of (D) CD18, (E) CD11a, (F) CD11b (G) CXCR2, (H) CD62L, (I) PSGL1 and (J) CD44 surface levels of WT and *Mrp14^-/-^* neutrophils [mean+SEM, *n=*3 mice per group, unpaired Student’s *t*-test]. (K) Schematic model of the adhesion rescue experiments in TNF-α stimulated WT and *Mrp14^-/-^* cremaster muscles by intra-arterial application of mutS100A8/A9 (aa exchange N70A + E79A). (L) Quantification of number of adherent WT and *Mrp14^-/-^* leukocytes mm^-2^ in the same vessel before and after mutS100A8/A9 i.v. injection [mean+SEM, *n=*4 mice per group, 4 (WT) and 4 (*Mrp14^-/-^*) vessels, 2way ANOVA, Sidak’s multiple comparison]. ns, not significant; *p≤0.05, **p≤0.01, ***p≤0.001.

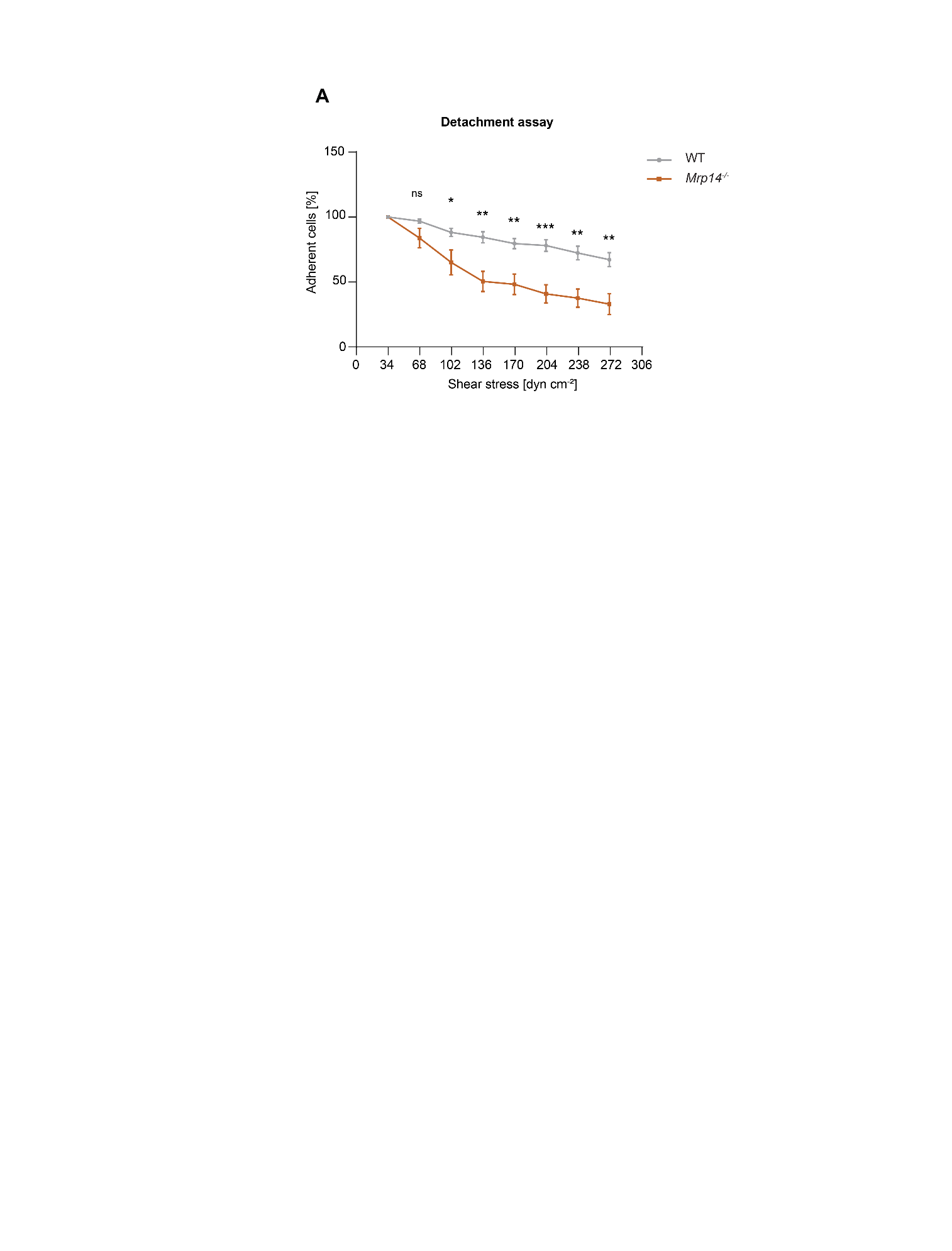

**SUPPLEMENTARY FIGURE 2**

**Figure S2: S100A8/A9 deficient cells are more susceptible to increasing shear stress compared to WT** **cells.** (A) Analysis of number of adherent WT and *Mrp14^-/-^* neutrophils under flow as percentage related to the initial number of adherent neutrophils in E-selectin, ICAM-1, and CXCL1 coated flow chambers at indicated shear stress levels. Shear stress was increased every 30sec. [mean+SEM, *n=*3 mice per group, 3 (WT) and 3 (*Mrp14^-/-^*) flow chambers, unpaired Student’s *t*-test]. ns, not significant; *p≤0.05, **p≤0.01, ***p≤0.001.

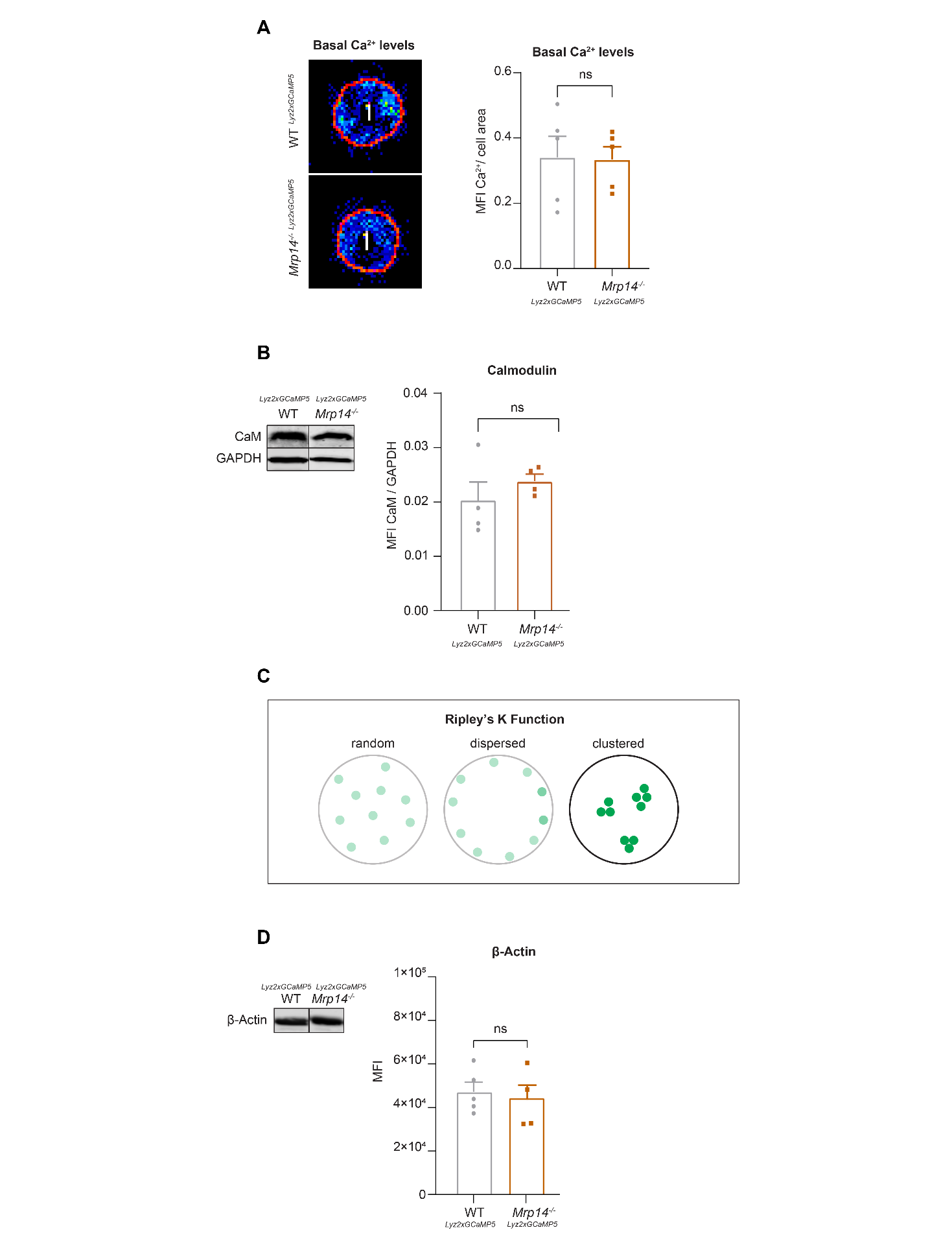
**SUPPLEMENTARY FIGURE 3**

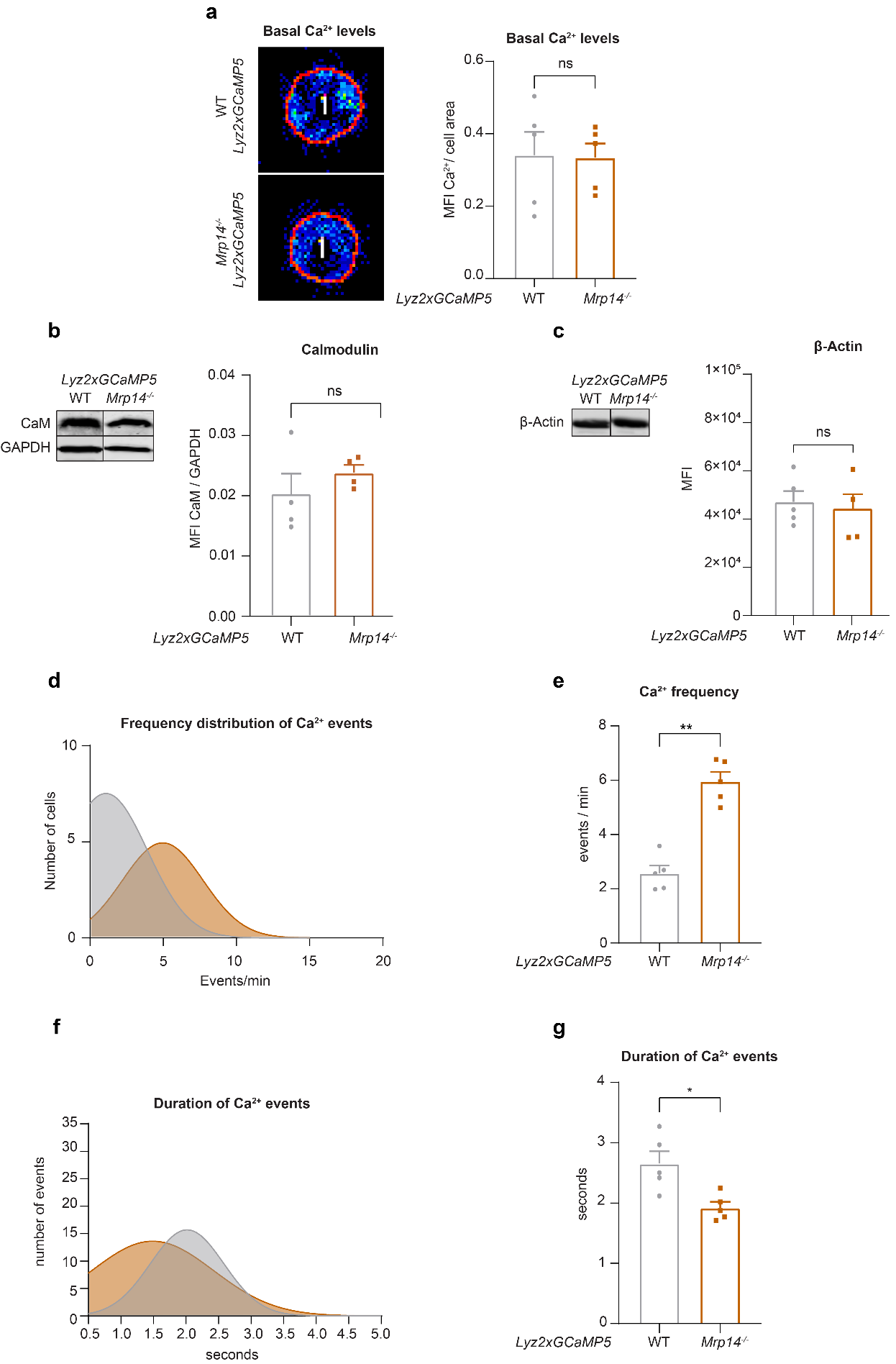

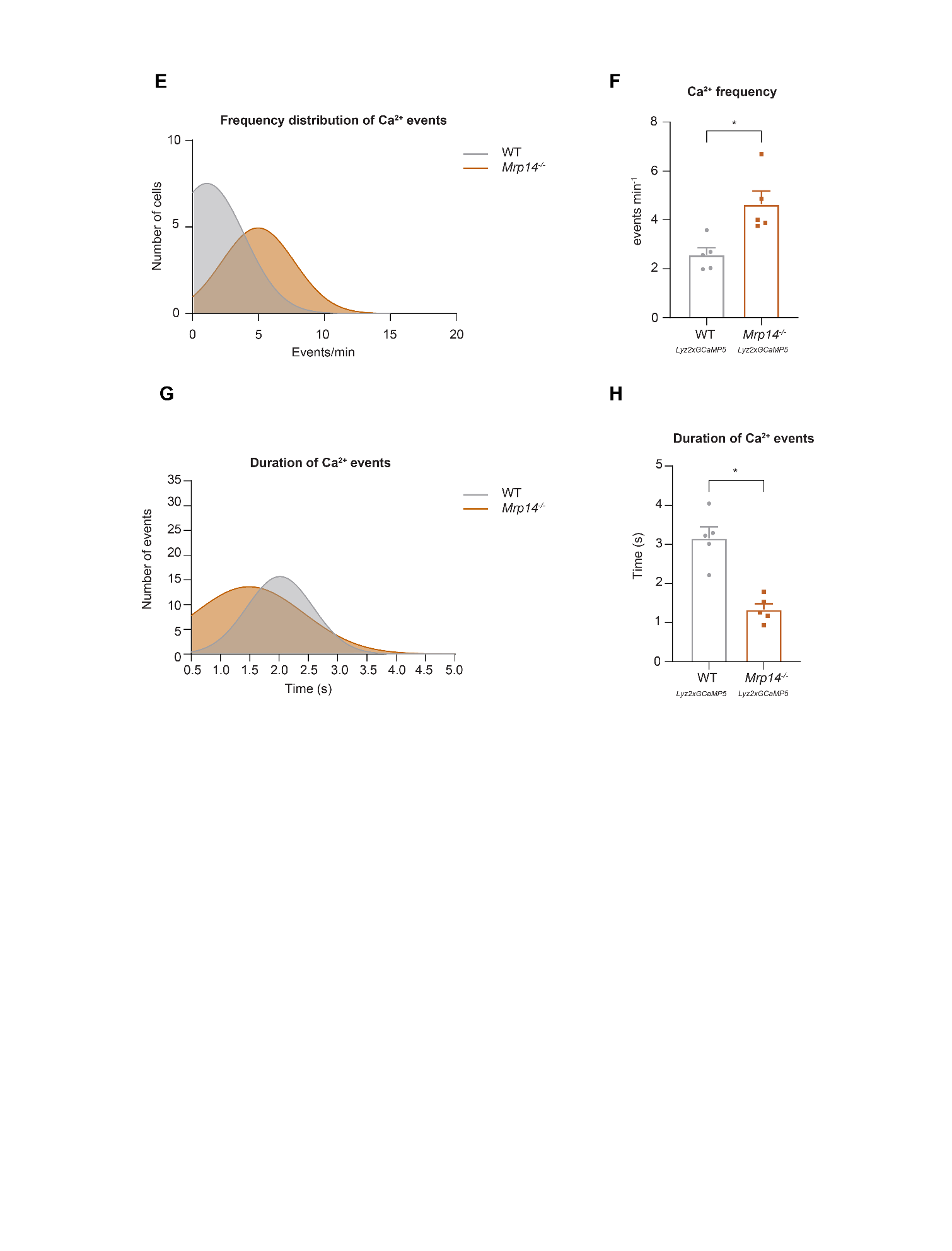

**Figure S3: S100A8/A9 deficient cells display higher frequencies but shorter duration of Ca^2+^ waves compared to WT cells.** (A) Analysis of basal Ca^2+^ levels normalized to the cell area of bone marrow derived WT *^Lyz2xGCaMP5^* and *Mrp14^-/-^ ^Lyz2xGCaMP5^* neutrophils seeded on poly-L-lysine coated slides [mean+SEM, *n=*234 (WT) and 192 (*Mrp14^-/-^*) neutrophils of 5 mice per group, paired Student’s *t*-test]. (B) Representative western blot images and quantification of total calmodulin levels normalized to GAPDH signal of WT *^Lyz2xGCaMP5^* and *Mrp14^-/-^ ^Lyz2xGCaMP5^* neutrophils [mean+SEM, representative western blot of *n=*4 mice per group, unpaired Student’s *t*-test]. (C) Schematic representation of the Ripley’s K used to determine spatial LFA-1 nanocluster correlation. (D) Representative western blot images and quantification of total actin levels (β-Actin) of WT *^Lyz2xGCaMP5^* and *Mrp14^-/-^ ^Lyz2xGCaMP5^* neutrophils [mean+SEM, representative western blot of *n≥*4 mice per group, unpaired Student’s *t*-test]. (E) Histogram of Ca^2+^ event frequency distribution and (F) quantification of Ca^2+^ event mean frequency in WT *^Lyz2xGCaMP5^* and *Mrp14^-/-^ ^Lyz2xGCaMP5^* neutrophils [mean+SEM, *n=*5 mice per group, paired Student’s *t*-test]. (G) Histogram of Ca^2+^ event duration and (H) quantification of average Ca^2+^ event duration in WT *^Lyz2xGCaMP5^* and *Mrp14^-/-^ ^Lyz2xGCaMP5^* neutrophils [mean+SEM, *n=*5 mice per group, paired Student’s *t*-test]. ns, not significant; *p≤0.05, **p≤0.01, ***p≤0.001.

**Table S1: Microvascular parameters in vivo**

Number of mice, number of vessels, vessel diameter, centerline velocity, wall shear rate and WBC of TNF-α stimulated WT and *Mrp14^-/-^* mice, as well as of WT and *Mrp14^-/-^* mice treated with mutS100A8/A9 without any prior stimulation (trauma model) and also of TNF-α stimulated WT and *Mrp14^-/-^* mice treated with mutS100A8/A9 (mean+SEM; unpaired student’s t-test).

|  | **Mice (n)** | **Venules (n)** | | **Diameter [µm]** | | | **Centerline velocity [µm s^-1^]** | **Wall shear rate [s^-1^]** | | **WBC**  **[µl^-1^]** |
| --- | --- | --- | --- | --- | --- | --- | --- | --- | --- | --- |
| **WT+ TNF-α** | 5 | | 21 | | 32.50+0.50 | 1130+50 | | 920+50 | 3580+450 | |
| ***Mrp14^-/-^*+ TNF-α** | 5 | | 20 | | 34+1.5 | 1320+50 | | 1010+50 | 3520+200 | |
|  |  | |  | | ns.  (p=0.6065) | ns.  (p=0.1534) | | ns.  (p=0.6091) | ns.  (p=0.9723) | |
| **WT + mutS100A8/A9** | 4 | | 4 | | 30+5 | 2030+300 | | 1800+420 | 5720+800 | |
| ***Mrp14^-/-^*+ mutS100A8/A9** | 3 | | 3 | | 30+2.5 | 1540+350 | | 1300+350 | 5460+650 | |
|  |  | |  | | ns.  (p=0.8359) | ns.  (p=0.3416) | | ns.  (p=0.3898) | ns.  (p=0.7979) | |
| **WT** **+ TNF-α + mutS100A8/A9** | 4 | | 18 | | 31+3 | 1330+330 | | 1450+400 | 4100+1000 | |
| ***Mrp14^-/-^*+ TNF-α + mutS100A8/A9** | 5 | | 24 | | 30+3 | 1700+130 | | 1500+300 | 3900+500 | |
|  |  | |  | | ns.  (p=0.6470) | ns.  (p=0.8341) | | ns.  (p=0.3947) | ns.  (p=0.6677) | |

**Table S2: Microvascular parameters ex vivo**

Number of mice, number of flow chambers, cells per FOV and WBC of ex vivo flow chamber assay (Mean+SEM, unpaired student’s t-test).

|  | **Mice (n)** | | **Flow chambers (n)** | | | **Cells FOV^-1^** | | **WBC**  **[µl^-1^]** |
| --- | --- | --- | --- | --- | --- | --- | --- | --- |
| **WT** | | 4 | | 8 | 39+5 | | 8630+1200 | |
| ***Mrp14^-/-^*** | | 4 | | 10 | 37+5 | | 8600+1200 | |
|  | |  | |  | ns.  (p=0.7332) | | ns.  (p=0.9772) | |

**Movie S1: Neutrophil functional crawling depends on S100A8/A9.** WT and *Mrp14^-/-^* neutrophils were seeded into E-selectin, ICAM-1 and CXCL1 coated flow chambers, allowed to settle down for 3min and then physiological shear stress (2dyne cm^−2^) was applied and crawling parameters were analyzed (representative time-lapse movies, frame interval 5sec, scale bar=10μm, time=min).

**Movie S2: S100A8/A9 is essential for LFA-1 nanocluster formation and turnover.** WT *^Lyz2xGCaMP5^* and *Mrp14^-/-Lyz2xGCaMP5^* neutrophils were seeded into E-selectin, ICAM-1 and CXCL1 coated flow chambers, allowed to settle down for 3min and then physiological shear stress (2dyne cm^−2^) was applied and number of LFA-1 nanoclusters was quantified at min 0-1, min 5-6 and min 9-10 of analysis (representative segmented and thresholded confocal microscopy movies of LFA-1 signals, scale bar=10μm).

**Movie S3: S100A8/A9 increases Ca^2+^ levels at the LFA-1 nanocluster sites.** WT *^Lyz2xGCaMP5^* and *Mrp14^-/-Lyz2xGCaMP5^* neutrophils were seeded into E-selectin, ICAM-1 and CXCL1 coated flow chambers, allowed to settle down for 3min and then physiological shear stress (2dyne cm^−2^) was applied and Ca^2+^ MFI was analyzed in the LFA-1 nanocluster areas at min 0-1, min 5-6 and min 9-10 of analysis [representative movies of Ca^2+^ oscillations in the LFA-1 segmented areas (“LFA-1 mask”), scale bar=10μm].

**Movie S4: S100A8/A9 induces F-actin polymerization.** WT *^Lyz2xGCaMP5^* and *Mrp14^-/-Lyz2xGCaMP5^* neutrophils were seeded into E-selectin, ICAM-1 and CXCL1 coated flow chambers, allowed to settle down for 3min and then physiological shear stress (2dyne cm^−2^) was applied and F-actin MFI was analyzed in the overall cell area at min 0-1, min 5-6 and min 9-10 of analysis [representative movies of F-actin intensity in the *Lyz2* segmented areas (“cell mask”), scale bar=10μm].
